# Experimental evolution and functional genetics identify KIC1 as a determinant of reduced artemisinin susceptibility in Bangladeshi *Plasmodium falciparum*

**DOI:** 10.64898/2026.07.17.738940

**Authors:** Maisha Khair Nima, Nirjhar Bhattacharyya, Saiful Arefeen Sazed, Ching Swe Phru, Mohammad Shafiul Alam, Michael T Ferdig, Angana Mukherjee

## Abstract

Bangladesh has greatly reduced malaria transmission, but persistent *Plasmodium falciparum* transmission in the Chittagong Hill Tracts (CHT), bordering Myanmar, remains a concern for elimination. Emerging artemisinin resistance could threaten remaining control efforts, making it important to identify determinants of reduced artemisinin susceptibility in CHT parasites. We combined long-term *in vitro* dihydroartemisinin (DHA) selection, whole-genome sequencing, functional genetics, and analysis of contemporaneous clinical isolates to define artemisinin-response variation in CHT *P. falciparum*. Starting with a 2018 patient-isolated, artemisinin-sensitive CHT clone, three independent cultures were exposed over 18 months to 27 cycles of stepwise DHA pressure up to 1600 nM. This generated CHT-R lines with elevated RSA survival relative to parental/sibling controls (4.0–6.5% versus <1%; 7–11-fold increase) and faster post-DHA recovery than CHT-S-sib (5.2 ± 0.3 versus 10.1 ± 0.57 days). Whole-genome sequencing identified convergent disruption of PF3D7_0606000, encoding KIC1 (Kelch13 Interacting Candidate1), a protein linked to the Kelch13-associated endocytic compartment; no *pfkelch13* mutations emerged. Independent DHA-selected lines acquired distinct stop-gain or frameshift mutations predicted to truncate KIC1, and targeted *pfkic1* disruption in parental CHT-S increased RSA survival and accelerated recovery, functionally validating KIC1 as a contributor to reduced artemisinin susceptibility. To link the *in vitro* selection findings to naturally circulating parasites, we tested whether *pfkic1* variation was associated with artemisinin response in an independent set of CHT isolates. We integrated newly generated RSA measurements and whole-genome variation from lab adapted field isolates all carrying wild-type *pfkelch13*, with patient clearance values from our CHT artemether-lumefantrine efficacy study. Among 22 isolates, RSA survival ranged from 0.00 to 6.44%, with 12 exceeding the 1% *in vitro* ART-R threshold and was associated with PC_50_, the time required to clear 50% of the initial parasite density, but not parasite clearance half-life (PCt_1/2_). In this field-isolate dataset, targeted *pfkic1* gene-score analysis and elastic-net modeling provided supportive evidence that natural variation at the *pfkic1* locus is associated with PC_50_ and RSA survival; an exploratory genome-wide gene-score scan nominated additional candidate loci for future study. Together, these findings identify KIC1 as a functionally validated determinant of reduced artemisinin susceptibility in a Bangladeshi parasite background and suggest that artemisinin-response variation in CHT parasites may involve perturbation of the Kelch13-associated endocytic pathway beyond canonical *pfkelch13* mutations.

## INTRODUCTION

Malaria caused by *Plasmodium falciparum* remains a major global health burden, and artemisinin-based combination therapies (ACTs) are the frontline treatment for uncomplicated falciparum malaria. The emergence and spread of artemisinin resistance (ART-R) therefore pose a major threat to malaria control and elimination efforts [1,2]. First identified in western Cambodia, ART-R subsequently spread across the Greater Mekong Subregion through both clonal expansion and independent emergence [3–7]. More recently, de novo emergence in eastern, southern and Horn of Africa has heightened concern that ART-R can arise in multiple geographic and genetic settings [8–14]. Clinically, ART-R is characterized by delayed parasite clearance (with slope half-lives; PCt_1/2_ >5h) [15], and *in vitro* it is commonly associated with elevated survival in the ring-stage survival assay (RSA) [16]. Mutations in *pfkelch13* are the best-established molecular markers of ART-R [17], discovered through *in vitro* evolution of ART-R in the lab from an isolate [17]. By 2023, the landscape of ART-R in Southeast Asia was dominated by the *pfkelch13* C580Y allele, which had largely displaced many earlier resistance-associated variants, although mutations such as F446I, R539T, P553L, and I543T continued to circulate in specific regional foci [18].

While *pfkelch13* mutations are central markers of ART-R, they do not fully explain the resistant phenotype. Increasing evidence from field, transcriptomic, and in vitro studies indicate that ART-R is a multigenic and genetic-background-dependent trait [19,20], that can either emerge on specific backgrounds or through Kelch13-independent determinants. In SEA and India *pfkelch13* WT parasites with prolonged clearance times and increased RSAs have been observed [5,21–24]. In Africa, reports from Tanzania, Mali, Kenya, and Uganda have identified delayed clearance, recurrent parasitemia, or treatment failure in infections lacking validated *pfkelch13* resistance mutations [25–30].

This polygenic, background-dependent trait of ART-R broadly converge on two main strategies, (i) limiting artemisinin activation, by reducing endocytosis; hemoglobin uptake and digestion, thereby reducing the heme-dependent activation of artemisinin. Kelch13 is involved in hemoglobin endocytosis and in Kelch13-mediated ART-R, mutations in *pfkelch13* impair Kelch13 function and disrupt endocytosis thereby limiting hemoglobin uptake and reducing artemisinin activation [31,32]. Kelch13 is a part of a multiprotein network at the cytostome [33] regulating hemoglobin uptake. This complex includes proteins such as AP-2u, UBP1, Eps15 and Kelch13 Interacting Candidates (KICs) [32–35]. Functional inactivation or modification of AP-2u, UBP1 and KIC7 reduced artemisinin susceptibility independent from Kelch13 [32,36,37]. (ii) Buffering artemisinin-induced cellular damage. Once activated by heme, artemisinins cause widespread cellular damage by oxidative stress, protein misfolding and DNA damage. Resistant parasites can tolerate drug induced damage through enhanced redox defense, proteostasis and DNA repair all of which have been supported by transcriptomic, population genomic, and functional studies in both *pfkelch13*-mutant and *pfkelch13*-wildtype backgrounds.

Bangladesh is an important but comparatively understudied setting for ART-R. Although the country achieved a 93% reduction in malaria cases between 2010 and 2020 and is actively pursuing malaria elimination by 2030 [38], approximately 18 million people remain at risk of malaria. *Plasmodium falciparum*, the major cause of severe malaria, accounts for most malaria infections in Bangladesh, with transmission concentrated in the forested Chittagong Hill Tracts (CHTs) of Bandarban, Rangamati, and Khagrachhari bordering Myanmar. In 2025, Bandarban and Rangamati accounted for most cases within the country, totaling 49% and 35% of cases respectively [39]. Artemether-Lumefantrine was introduced as the official first-line ACT therapy for uncomplicated *P. falciparum* malaria in 2005 [40] and no reports of validated ART-R associated *pfkelch13* mutations have been detected in the CHTs nor Bangladesh. In this pre-elimination phase, increasing concern exists that continued ACT use together with cross-border movement of resistant parasites from neighboring Myanmar could facilitate the emergence or introduction of ART-R into Bangladesh and threaten malaria elimination efforts [41].

In 2018–2019, we assessed ACT susceptibility in *P. falciparum* monoinfected malaria patients in clinical parasite clearance assays after artemether-lumefantrine treatment in the CHTs in Bangladesh [42]. All isolates exhibited PCt_1/2_ <5h [42], demonstrating sensitive clinical phenotypes per WHO definition. However, the median time to clear half of the initial parasite load (PC_50_) was 5.6 h (range: 1.5–9.6 h), with 20% of patients exhibiting a median of 8 h [42]. Additionally, we also detected and reported quantifiable *in vitro* resistance by RSA that was independent of Kelch13 substitutions [42,43]. These findings raised the possibility that CHT parasites may harbor variation in artemisinin response that is not captured by conventional delayed clearance (half-life) or validated *pfkelch* 13 mutations.

To identify genetic changes that can emerge under artemisinin pressure in CHT parasites, we integrated long-term *in vitro* DHA selection with phenotypic and genomic analysis of natural CHT isolates. Stepwise DHA selection of a recently adapted, artemisinin-sensitive CHT isolate generated independent lines with elevated RSA survival and recurrent predicted loss-of-function mutations in PF3D7_0606000 (*pfkic1*), encoding KIC1, the top-ranked Kelch13-interaction candidate. Targeted disruption of *pfkic1* in the parental background was sufficient to increase RSA survival and post-DHA recovery, functionally validating KIC1 as a contributor to reduced artemisinin susceptibility. We then used contemporaneous CHT clinical isolates to ask whether KIC1 was also implicated in local artemisinin-response variation. In these isolates, *pfkic1* score was nominally associated with higher PC_50_ and elevated RSA survival, and an unbiased genome-wide scan identified additional candidates in vesicle-trafficking and proteostasis pathways. Together, these data identify KIC1 as a Kelch13-independent determinant of reduced artemisinin susceptibility in a Bangladeshi parasite background and suggest that local artemisinin-response variation may involve broader perturbation of genes linked to Kelch13-associated endocytosis.

## RESULTS

### *In vitro* DHA Selections of a CHT Isolate Generate Elevated Ring-Stage Survival and Faster Post-DHA Recovery

To identify genetic variants arising under artemisinin drug pressure that may contribute to partial ART-R, we subjected a Bangladeshi *Plasmodium falciparum* patient isolate to intermittent *in vitro* selection with progressively increasing concentrations of dihydroartemisinin (DHA), the active metabolite of artemisinin (**Fig. 1**).The parental line, CHT-S, was a laboratory-adapted, monogenomic isolate (I-001) collected in 2018 from Bandarban in the CHT of Bangladesh. The patient from whom this isolate was obtained exhibited rapid parasite clearance following artemether–lumefantrine treatment (PC₅₀ = 0.015 h; PCt_1/2_= 3.3 h), and the isolate displayed a DHA-sensitive *in vitro* RSA phenotype after adaptation (0.66 ± 0.13% survival) with a wild-type *kelch13* sequence [42]. *In vitro* selection was initiated using three independent flasks of CHT-S. Two flasks were exposed to intermittent 24 h pulses of 40 nM DHA, while a third flask was exposed to 48 h pulses of 20 nM DHA. Following each DHA exposure, parasites were cultured drug-free until recovery to 1% parasitemia; the interval from the start of the DHA pulse to recovery defined one selection cycle. A sibling control line was maintained in parallel without DHA exposure to control for mutations arising from prolonged *in vitro* culture (**Fig. 1A**). DHA concentrations were increased stepwise across successive selection cycles (40, 60, 80, 100, 200, 400, 600, 800, 1200, and ultimately 1600 nM; 800× EC₅₀). During early stages of selection, if parasites recovered within 12–14 days, the same DHA concentration was reapplied in the subsequent cycle; if recovery required more than 12 days, the next higher DHA concentration was applied. Recovery times between selection cycles are shown in **Fig. 1B**, with a maximum recovery time of 21 days observed across all selections. By approximately the 20th selection cycle, corresponding to 800 nM DHA, all three selections consistently recovered to 1% parasitemia in less than 10 days and maintained similar recovery kinetics at higher DHA concentrations (**Fig. 1B**). After ∼18 months of selection, three stable, independently selected lines, CHT-R-24-1, CHT-R-24-2, and CHT-R-48 were established (**Fig. 1A**). We performed Ring- Stage Survival Assays (RSA) on the selected lines, alongside the parent (CHT-S) and the long-term cultured sibling control (CHT-S-sib) at selected selection points (see **Fig 1D**, for selected points). Synchronized 0–3 h ring-stage parasites were exposed to 700 nM DHA for 6 h, and survival was measured 66 h after drug removal by comparing DHA-treated parasites with matched DMSO-treated controls. Selected lines revealed a sustained increase in survival rates in DHA selected lines compared to parental CHT-S and CHT-S-sib (**Fig 1C**). CHT-R-24 lines exhibited significant increase in survival rates with a mean % survival of 4- 5.7% at the 1600nM endpoint. Similarly, CHT-R-48 lines demonstrated a slightly higher survival rates than CHT-R-24 lines, with a % survival range of 5.2-6.5% at the 1600nM endpoint. This represents a 7-11X increase compared to the parental and sibling with % survival of 0.57% ± 0.117 (mean ± S.E.M). Statistical analysis using one-way ANOVA with Dunnett’s multiple-comparison test showed significantly increased RSA survival in the DHA-selected lines compared with CHT-S-sib, with significant comparisons indicated in **Fig. 1C** (*P* < 0.05). We next assessed the recovery of these selected lines following 700 nM DHA pulse to 0-3h rings for 6 h by measuring the time required to reach 0.5% parasitemia from the start of the assay (**Fig. 1D**). The CHT-S-sib required 10.1 ± 0.57 days (mean ± SEM) to recover, whereas, the drug-selected lines CHT-R-24-1, CHT-R-24-2, and CHT-R-48 recovered within approximately 8–10 days, similar to the parent line, through selection cycle ∼21 (corresponding to ∼800 nM DHA selection pressure). Increasing selection dosages corresponded to improved recovery times after RSA. Toward the end of the selection process (27th selection cycle) at 1600 nM, all CHT-R lines required significantly fewer days for recovery, with a mean recovery time of 5.2 ± 0.3 days (mean ± S.E.M), compared to the parental CHT-S strain (**Fig. 1D**). RSA survival was inversely associated with post-DHA recovery time across selected lines and selection stages (Spearman ρ = −0.912, *P* < 0.0001, n = 17; linear regression, *R*² = 0.796, *P* < 0.0001) (**Fig. 1E**).

**Fig. 1.**
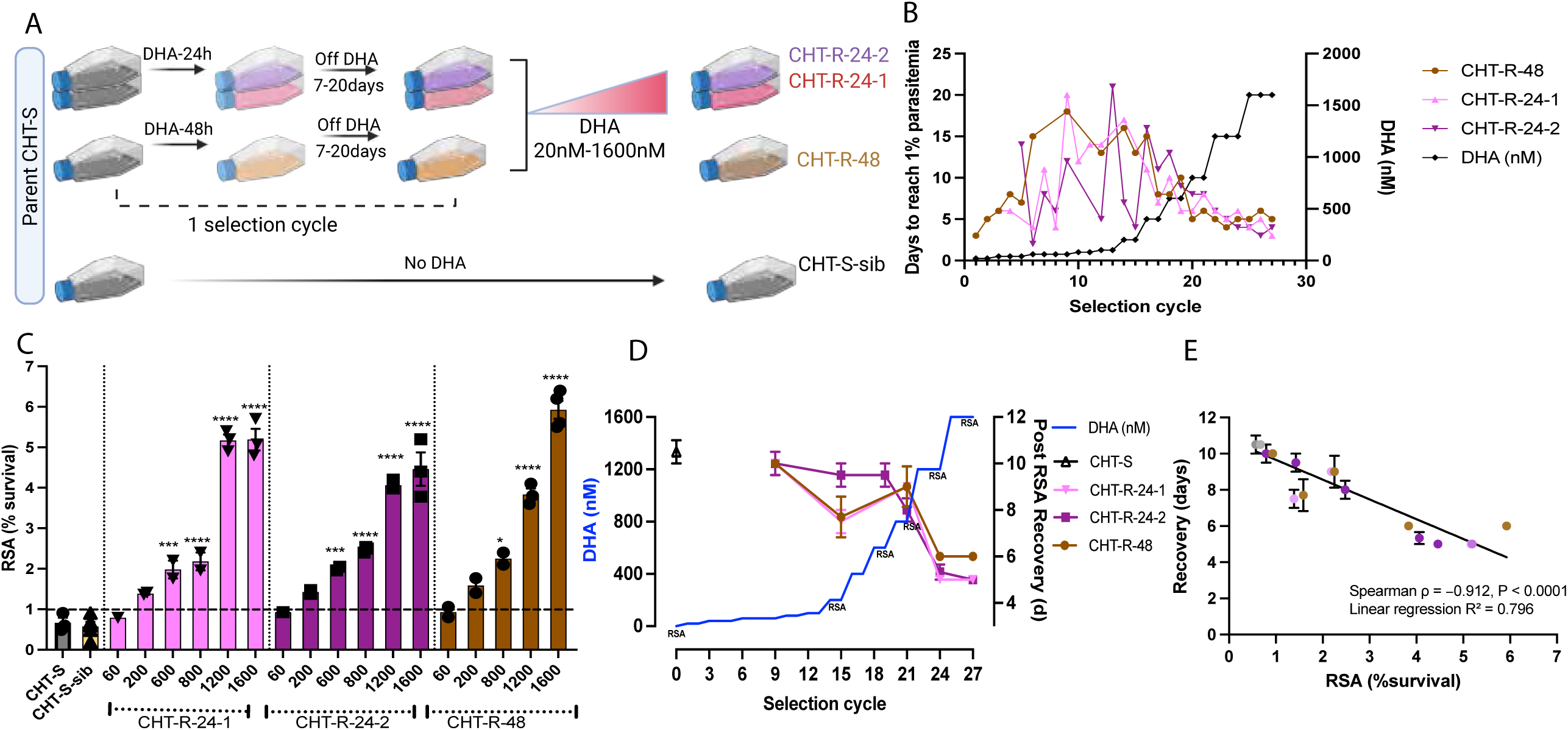
*In vitro* DHA selection of a Bangladeshi CHT isolate (CHT-S) generates elevated RSA survival and faster post-DHA recovery. **(A)** Schematic of the DHA selection strategy. Three independent CHT-S cultures were exposed to intermittent DHA pressure: two with 24 h pulses and one with 48 h pulses. After each DHA exposure, parasites were cultured drug-free until recovery to 1% parasitemia, defining one selection cycle. DHA was increased stepwise to 1600 nM, while CHT-S-sib was maintained without DHA. **(B)** Recovery time during stepwise DHA selection, defined as days from DHA exposure to recovery at 1% parasitemia. Recovery times are shown across selection cycles for each independently selected line as DHA pressure increases over time. **(C)** RSA survival of parental CHT-S, sibling CHT-S-sib, and DHA-selected lines after exposure of synchronized 0–3 h rings to 700 nM DHA for 6 h. Survival was measured 66 h after drug removal relative to matched DMSO controls. The dashed line indicates the 1% RSA threshold. Data are shown as mean ± SEM. RSA survival was compared across parental, sibling, and DHA-selected lines using one-way ANOVA with Dunnett’s multiple-comparison test; selected lines with significantly increased RSA survival compared to CHT-S-sib are indicated. *p=<0.05; ***p=<0.001; ****p=<0.0001. **(D)** Recovery Assay. Post-DHA recovery of parasites after 0-3h parasites were exposed to 700nM DHA for 6h, measured as days required to return to 0.5% parasitemia. Right Y-axis shows recovery time (days) to reach 0.5% parasitemia. Data are shown as mean ± SEM. Left Y-axis shows the DHA concentration applied for selections. Selection points where RSAs (**1E**) and Recovery Assays were performed are indicated as “RSA”. **(E)** Relationship between RSA survival and post-DHA recovery time across DHA-selected lines and selection stages. RSA survival and recovery time were negatively correlated, indicating that lines with higher RSA survival recovered more rapidly after a 700 nM DHA pulse for 6 h (Spearman ρ = −0.912, *P* < 0.0001; linear regression, *R*² = 0.796, *P* < 0.0001).

### Genome sequencing of DHA selected lines and candidate genes associated with ART-R

We performed whole-genome sequencing to identify single nucleotide polymorphisms (SNPs) in the artemisinin selected lines and detected multiple SNPs relative to both CHT-S and CHT-S-sib. Whole genome sequences were obtained at an average of 214X coverage with 39698092.17(87%) reads mapped on the genome (**Table S1**). Variants of each 1600nM artemisinin selected line were compared to the parent and sibling lines after excluding SNPs from highly variable multi-gene families. SNPs shared between each 1600nM line and either the parent or the sibling clone were excluded. The CHT-S-sib had more sporadic SNPs compared to the DHA selected lines possibly since the DHA selection was hard and targeted to induce mutations in genes that can confer resistance or survival; whereas culturing condition was more of an untargeted selection pressure. The overall distribution of annotated SNP classes was broadly similar across the parental, sibling, and DHA-selected lines, with missense and intergenic variants representing the largest categories, followed by synonymous and intronic variants (**Fig. S1**). INDEL annotations were similarly dominated by intergenic and intronic variants, with a smaller fraction classified as frameshift or in-frame disruptive events. After removing variants shared with the CHT-S-sib, the remaining DHA-selected-line variants were enriched for intergenic changes, while genic variants comprised approximately 30–48% of the remaining calls. We focused subsequent analysis on high-confidence genic variants; nonsynonymous and loss-of-function mutations absent from both the parental and sibling controls. SNP output per filtering step for each selected line is in **Table S2**. Mutations in some candidate genes were unique to one selected line, while others were shared (**Table S3**). Our analysis identified seventeen unique SNPs shared across six genes in all three 1600nM selected lines (**Table 1**). Different mutations in PF3D7_0606000 encoding the protein KIC1 (Kelch13 Interaction Candidate 1), the top-ranked Kelch13-interaction candidate identified in the Kelch13 BioID/proximity-labeling dataset [32] were identified in all the three selected lines. CHT-R-24-1 acquired a Ser186* stop-gain mutation, CHT-R-24-2 harbored a frameshift insertion (874Ins) that resulted in a premature stop codon, and CHT-R-48 developed a Ser177* stop-gain mutation. The recurrent emergence of unique *pfkic1* mutations across all selections strongly implicates this gene in ART-R in the CHT-S genetic background.

**Table 1:** Genes recurrently mutated across the CHT-R-24-1, CHT-R-24-2, and CHT-R-48 parasite lines with non-identical variant profiles. Genes were included when at least one protein-altering variant was detected in each of the three lines. “Present” indicates detection of the specified variant in the corresponding line, whereas “—” indicates that the variant was not detected. Asterisks (*) indicate premature termination codons produced by stop-gained variants.

| Gene ID<br>(PlasmoDB) | Gene name | Chr:Pos | Ref<br>>Alt | Effect | Amino acid<br>change | CHT-R-24-1 | CHT-R-24-2 | CHT-R-48 |
| --- | --- | --- | --- | --- | --- | --- | --- | --- |
| PF3D7_1227500 | CYC2/SOC<br>2 | Pf3D7_12_v3:<br>1114059 | T>C | missense | Tyr2293Cys | present | present | present |
|  |  | Pf3D7_12_v3:<br>1117183 | G>A | missense | Ala1304Val | — | — | present |
| PF3D7_0405300 | LISP2 | Pf3D7_04_v3:<br>285494 | T>G | missense | Asp935Ala | present | present | — |
|  |  | Pf3D7_04_v3:<br>285425 | G>A | missense | Ala958Val | — | — | present |
| PF3D7_0220000 | LSA3 | Pf3D7_02_v3:<br>798072 | A>T | missense | Glu388Val | — | — | present |
|  |  | Pf3D7_02_v3:<br>798078 | G>T | missense | Ser390Ile | — | present | — |
|  |  | Pf3D7_02_v3:<br>798102 | T>G | missense | Ile398Ser | — | — | present |
|  |  | Pf3D7_02_v3:<br>801565 | T>A | missense | Ser1552Arg | present | present | present |
| PF3D7_0423600 |  | Pf3D7_04_v3:<br>1064402 | T>C | missense | Thr743Ala | present | present | — |
|  |  | Pf3D7_04_v3:<br>1064339 | T>C | missense | Thr764Ala | — | — | present |
| PF3D7_0606000 | KIC1 | Pf3D7_06_v3:<br>253247 | G>T | stop<br>gained | Ser186* | present | — | — |
|  |  | Pf3D7_06_v3:<br>251183 | C>T | missense | Ser874Asn | — | present | — |
|  |  | Pf3D7_06_v3:<br>253274 | G>T | stop<br>gained | Ser177* | — | — | present |
| PF3D7_0628800 | GATB | Pf3D7_06_v3:<br>1187818 | A>G | missense | Val276Ala | present | present | present |
|  |  | Pf3D7_06_v3:<br>1187819 | C>A | missense | Val276Leu | present | present | present |
|  |  | Pf3D7_06_v3:<br>1187834 | C>A | missense | Val271Leu | present | present | present |
|  |  | Pf3D7_06_v3:<br>1187848 | T>G | missense | Glu266Ala | present | — | — |
|  |  | Pf3D7_06_v3:<br>1187848 | T>A | missense | Glu266Val | — | present | present |

### Genetic Disruption of *pfkic1* is tolerated in CHT-S but not in *pfkelch13*-mutant backgrounds

To functionally evaluate the role of KIC1 in artemisinin susceptibility, we genetically disrupted *pfkic1* in the parental CHT-S line. Because all independently selected lines acquired premature stop codons in *pfkic1*, we reasoned that the gene is likely non-essential and therefore we attempted gene disruption using a targeted selection-linked integration strategy (SLI-TGD) [32] and **Fig. S2A**. We generated three independent *pfkic1*-disrupted lines (*kic1*-TGD-1, *kic1*-TGD-2, and *kic1*-TGD-3) in the CHT-S background. PCR analysis confirmed modification of the *pfkic1* locus in all three lines, with bands corresponding to the integrated plasmid at 5 ‘and 3’ junctions (**Fig. S2B**) and the absence of the original WT allele band in all three TGD lines, consistent with successful disruption of the original endogenous *pfkic1* locus (**Fig. S2B**). These results confirm that *pfkic1* is not essential in the asexual stages in the CHT-S background. We also attempted to disrupt *pfkic1* in two *pfkelch13* edited CHT-S (CHT-S^C580Y^ and CHT-S^F446I^) lines [43] that demonstrated RSAs of 4.47% ± 0.4% and 1.34% ± 0.33% (mean ± SEM) respectively [43] to see the effect of artemisinin susceptibility in these double transgenics. Despite repeated (six) attempts, we failed to generate *pfkic1* disruptions in the presence of Kelch13 substitutions. This may suggest that *pfkic1* disruption in *kelch13*-mutated parasites is not viable in the CHT-S genetic background, highlighting a potential genetic dependency or fitness cost associated with *kelch13* mutations when combined with *pfkic1* disruption.

### Validation of the role of KIC1 in artemisinin susceptibility

To assess the effect of *pfkic1* disruption in artemisinin susceptibility, we performed RSAs and post-DHA recovery assays. Consistent with their artemisinin-sensitive phenotype, both the CHT-S and the CHT-S-sib exhibited RSA survival rates well below the 1% threshold for ART-R (parent: 0.66 ± 0.13%; sibling: 0.91 ± 0.10%; mean ± SEM) (**Fig. 2A**). In contrast, all three *kic1*-TGD lines showed markedly elevated RSA survival. *kic1*-TGD-1 exhibited 8.886 ± 1.407% survival, corresponding to a 13.4-fold increase relative to the parent and a 10.6-fold increase relative to the sibling. Similarly, *kic1*-TGD-2 showed 10.58 ± 0.4591% survival (15.9-fold higher than the parent; 11.6-fold higher than the sibling), and *kic1*-TGD-3 showed 10.04 ± 1.387% survival (15.1-fold higher than the parent; 11.0-fold higher than the sibling). All three disrupted lines differed significantly from the sibling CHT-S line (Welch’s *t*-test, *p* < 0.01) (**Fig. 2A**). We next examined whether the elevated RSA survival of the *kic1*-TGD lines translated into enhanced recovery following DHA exposure. Consistent with their low RSA survival, both CHT-S and CHT-S-sib recovered poorly, with parasitemia remaining below 0.5% throughout the 192h observation period (**Fig. 2B**). In contrast, all three *kic1*-TGD lines exhibited rapid post-DHA expansion, with parasitemia increasing steadily after 96–120 h and reaching approximately 2–2.5% by 168 h (**Fig. 2B**).

**Fig. 2.**
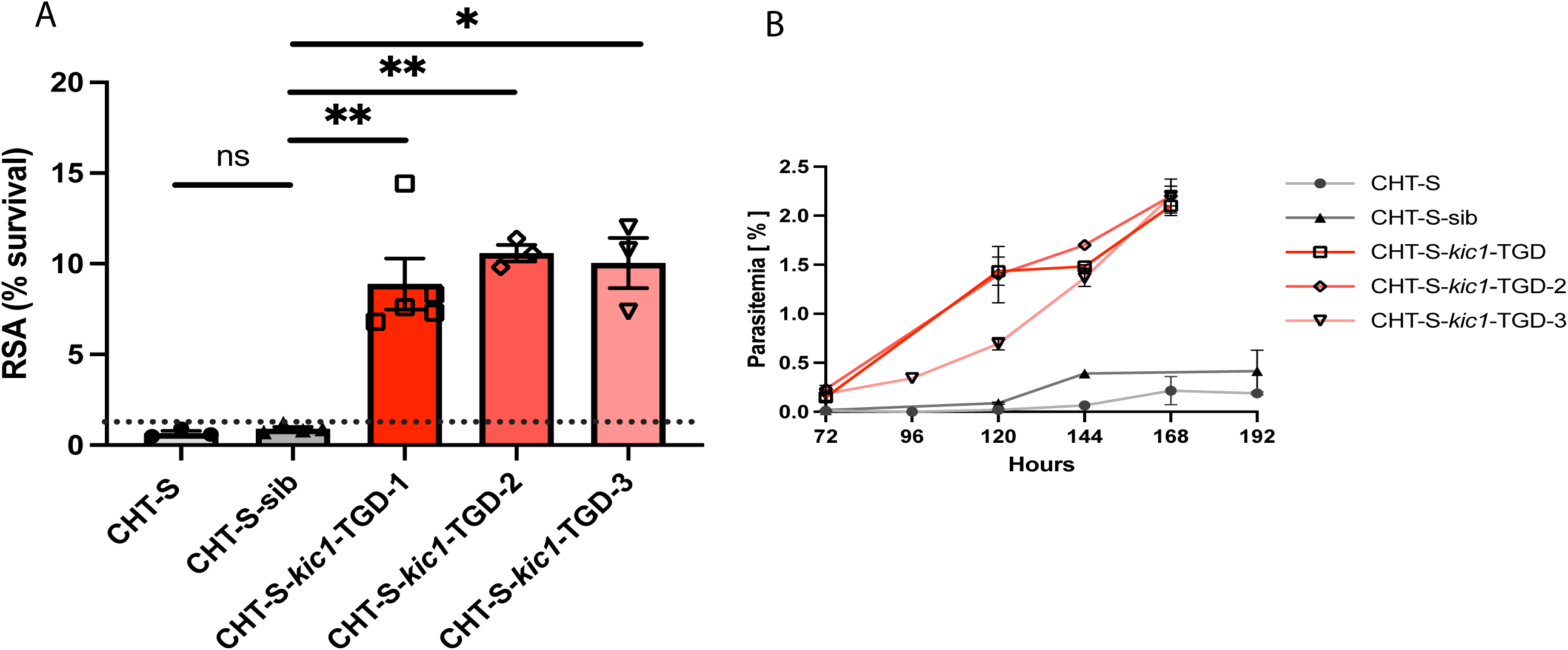
Disruption of *pfkic1* increases RSA survival and post-DHA recovery in the CHT-S background. **(A)** Mean ± SEM values from 3-5 independent replicates of RSA survival of parental CHT-S, sibling control CHT-S-sib, and three independent *kic1*-TGD lines. Synchronized 0–3 h rings were exposed to 700 nM DHA for 6 h, and survival was measured 66 h after drug removal relative to matched DMSO controls. The dashed line indicates the 1% RSA survival threshold. Welch’s t test was performed between CHT-S-sib and all other lines. p=0.1944 (CHT-S); p=0.0047 (*kic1*-TGD-1); p=0.0015 (*kic1*-TGD-2); p=0.0218 ((*kic1*-TGD-3). **(B)** Post-DHA recovery of CHT-S, CHT-S-sib, and *kic1*-TGD lines after 700 nM DHA exposure to 0-3h rings for 6 h. Parasitemia was monitored over 192 h after drug exposure.

### RSA survival in contemporaneous CHT clinical isolates is associated with PC_50_ but not parasite clearance half-life

To assess artemisinin susceptibility in natural parasites from the CHTs, we built on our previously published parasite clearance study conducted at Sadar Hospital in Bandarban in 2018–2019 [42]. The CHTs, comprising Bandarban, Rangamati, and Khagrachhari districts, account for the majority of *P. falciparum* malaria in Bangladesh, and Bandarban was the site of our clinical sampling. In that study, 41 patients with *P. falciparum* monoinfection were enrolled, and parasite clearance following artemether–lumefantrine treatment was estimated over 72 h in 24 patients [42]. Consistent with prior studies from Bangladesh [44,45], sequencing of *pfkelch13* from these isolates did not identify validated ART-R-associated propeller-domain mutations [42]. ACT treatment remained clinically efficacious, as no patients had detectable parasitemia on day 3 or a PCt_1/2_ ≥5 h [42] and **Table 2**, the WHO-associated criteria used to define delayed parasite clearance in the Greater Mekong Subregion [46]. Although all infections met the WHO criteria for rapid parasite clearance, the median time to clear 50% of the initial parasite density (PC_50_) was 3.5 h (range, 0.015–7.6 h), with 9 of 24 patients exhibiting PC_50_ values >5 h [42] and **Table 2**.

**Table 2:** Patient parasite clearance half-lives (PC_50_ and PCt_1/2_) [42] and *in vitro* RSA survival values of the corresponding isolates (n=3, except O-024, n=4). *Whole genome sequences performed on these lab-adapted isolates

| Strains | Patient Parasite Clearance Times (h) |  | RSA (% survival) |  |
| --- | --- | --- | --- | --- |
| | $PC_{t_{1/2}}$ (h) | $PC_{50}$ (h) | Mean | SEM |
| I-001* | 3.3 | 0.015 | 0.6636 | 0.126 |
| I-002 | NA | 3.56 | 2.563 | 0.2 |
| I-003* | 3.39 | 5.967 | 2.007 | 0.1195 |
| I-004* | 2.25 | 1.6 | 0 | 0 |
| I-005 | 2.49 | 3.82 | NA | NA |
| I-008* | 1.42 | 2.71 | 1.536 | 0.1976 |
| I-010 | 1.42 | 2.01 | NA | NA |
| I-011* | 2.36 | 5.35 | 0.6058 | 0.3178 |
| I-013 | 1.56 | 7.98 | NA | NA |
| I-016* | 1.77 | 10.25 | 4.635 | 0.5042 |
| I-017 | 1.83 | 4.51 | NA | NA |
| I-018* | 1.86 | 3.42 | 0.7467 | 0.03528 |
| I-019* | 2.65 | 2.02 | 0.4633 | 0.08819 |
| I-020* | 2.14 | 7.66 | 1.76 | 0.3378 |
| I-021 | 1.22 | 2.83 | NA | NA |
| I-025 | 2.43 | 3.39 | NA | NA |
| I-026* | 1.62 | 3.25 | 0.6533 | 0.06119 |
| I-027* | 2.89 | 3.23 | 0.5593 | 0.1399 |
| I-029* | 2.39 | 7.68 | 6.443 | 0.3096 |
| I-030 | 2.04 | 8.19 | NA | NA |
| I-031 | 1.52 | 3.41 | NA | NA |
| I-033* | 2.42 | 8.48 | 1.89 | 0.3074 |
| I-035 | 3.25 | 1.25 | NA | NA |
| I-036 | 2.69 | 9.42 | NA |  |
| I-039* | 2.62 | 13.14 | 4.892 | 0.1202 |
| I-040* | 2.17 | 3.14 | 0.7467 | 0.05812 |
| I-041* | 2.47 | 4.82 | 1.359 | 0.265 |
| O-009* | NA | NA | 0.42 | 0.07572 |
| O-014 | NA | NA | 2.127 | 0.3634 |
| O-024* | NA | NA | 1.73 | 0.3957 |
| O-032* | NA | NA | 0.783 | 0.06358 |
| O-038* | NA | NA | 1.372 | 0.0955 |

To further evaluate whether these parasites exhibited reduced artemisinin susceptibility *in vitro*, 17 cryopreserved isolates from this study were shipped to Notre Dame for genomic and *in vitro* artemisinin phenotypic analysis. In addition, five isolates collected from outpatient cases in 2018 without parasite clearance data (O-014, O-024, O-027, O-032, and O-038) were also cryopreserved and shipped to Notre Dame for culture adaptation and included for *in vitro* analysis. In total, we short-term adapted 22 *P. falciparum* isolates from Bandarban for phenotypic and genetic profiling and measured their artemisinin susceptibility using the ring-stage survival assay (RSA). Survival was benchmarked against the NF54 (*kelch13* WT) and edited, isogenic NF54 K13^C580Y^ (**Fig. 3A**). RSA values of these comparators aligned with published values [47]. RSA survival among the 22 CHT isolates ranged from 0.00 to 6.44%, with a median survival of 1.37% (IQR, 0.64–2.04%). Twelve of 22 isolates (54.5%) exceeded the 1% RSA threshold for *in vitro* ART-R (**Fig. 3A**). We next asked whether *in vitro* RSA survival was associated with clinical parasite clearance parameters. RSA survival was strongly correlated with PC_50_, the time required to clear 50% of the initial parasite density (Spearman ρ = 0.77, P = 0.004; **Fig. 3B**). In contrast, RSA survival was not associated with parasite clearance half-life (Spearman ρ = −0.02, P = 0.95; **Fig. 3C**). Thus, in this CHT cohort, elevated ring-stage survival was associated with early parasite clearance dynamics but not with the conventional clearance half-life metric.

**Fig. 3.**
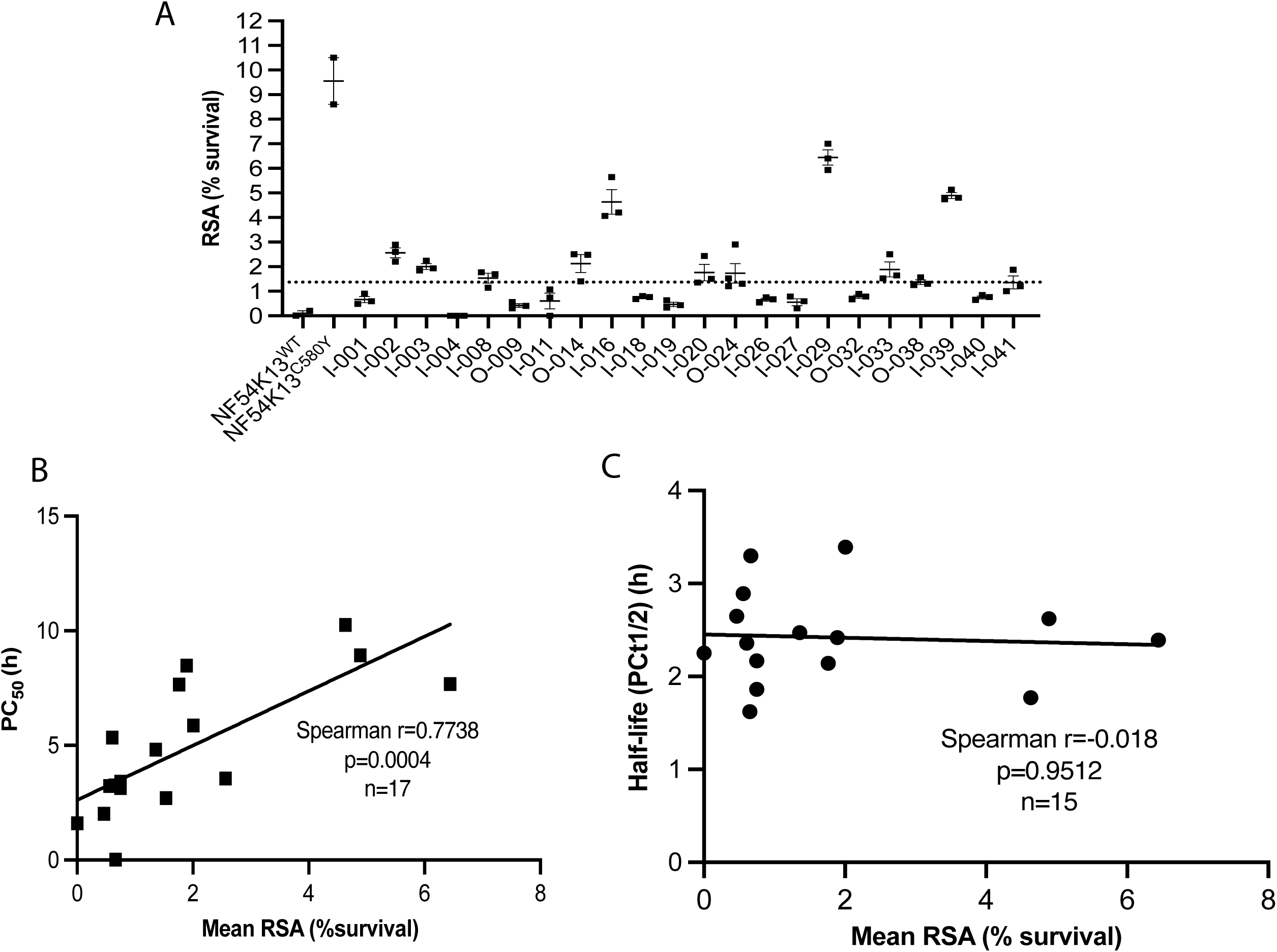
CHT clinical isolates show heterogeneous RSA survival associated with PC_50_ but not parasite clearance half-life. **(A)** RSA survival among 22 culture-adapted *P. falciparum* isolates from Bandarban, Bangladesh. NF54 Kelch13 WT and isogenic NF54 Kelch13-C580Y were included as DHA-sensitive and ART-R comparator controls, respectively. Synchronized 0–3 h rings were exposed to 700 nM DHA for 6 h, and survival was measured 66 h after drug removal relative to matched DMSO controls. Data are shown as mean ± SEM RSA survival from individual isolates. The dashed line indicates the median value of the RSAs from all isolates. **(B)** Association between mean RSA survival and PC_50_, defined as the time required to clear 50% of the initial parasite density after artemether-lumefantrine treatment. RSA survival was positively correlated with PC_50_. Spearman ρ = 0.7738, *P* = 0.0004, n = 17. **(C)** Association between mean RSA survival and parasite clearance half-life. RSA survival was not associated with parasite clearance half-life. Spearman ρ = −0.018, *P* = 0.9512, n = 15

### Rare-variant gene-score analysis identifies nominal associations between the *pfkic1* locus score and PC_50_/RSA phenotypes in the CHT clinical isolates

To test whether *pfkic1* variation was associated with artemisinin-response phenotypes in CHT clinical isolates, we constructed a rare-variant, Madsen–Browning-weighted gene score for *pfkic1* and tested it against PC_50_ and *in vitro* RSA survival. Because potentially relevant *pfkic1* alleles were individually rare, nonsynonymous and proximal variants at the *pfkic1* locus were collapsed into a single weighted score that upweighted rarer alleles [48]. PC_50_, rather than PCt_1/2_, was prioritized because it captured early parasite-clearance variation associated with RSA survival in this cohort (**Fig. 3C**). Seven *pfkic1* variants were retained for the PC_50_ analysis (n = 15; **Table S6**), with MAF-derived weights ranging from 2.94 to 4.01 (**Table S5**). OLS regression of continuous PC_50_ against the *pfkic1* gene score showed a nominal positive association (β = 0.167, *P* = 0.019, *R*² = 0.357) (**Fig. 4A**). Pearson correlation was consistent with this association (*r* = 0.598, *P* = 0.019), and Spearman rank correlation also supported a positive relationship (ρ = 0.655, *P* = 0.008). As a supportive group-based analysis, we stratified isolates by PC50 ≥5 h versus PC50 <5 h to compare parasites with slower versus faster early clearance kinetics; this grouping was not intended to define WHO clinical artemisinin resistance. The *pfkic1* gene score differed between the two PC50 groups by Mann–Whitney/Wilcoxon rank-sum test (*P* = 0.019) (**Fig. S3A**). We next tested the same *pfkic1* locus score against *in vitro* RSA survival. Among 22 isolates, 12 had RSA survival ≥1% and 10 had RSA survival <1% (**Table 2 and Table S4**). OLS regression of continuous RSA survival against the *pfkic1* gene score showed a nominal positive association (β = 0.067, *P* = 0.025, *R*² = 0.250) (**Fig. 4B**), with the score accounting for approximately 25% of the variation in RSA survival across 20 isolates (two, I-002 and O-014 were excluded as they failed QC) in this exploratory model. Pearson correlation was consistent with this relationship (*r* = 0.500, *P* = 0.025). However, group-level comparison of *pfkic1* scores between isolates with RSA survival ≥1% and <1% did not reach nominal significance (**Fig. S3B**).

**Fig. 4.**
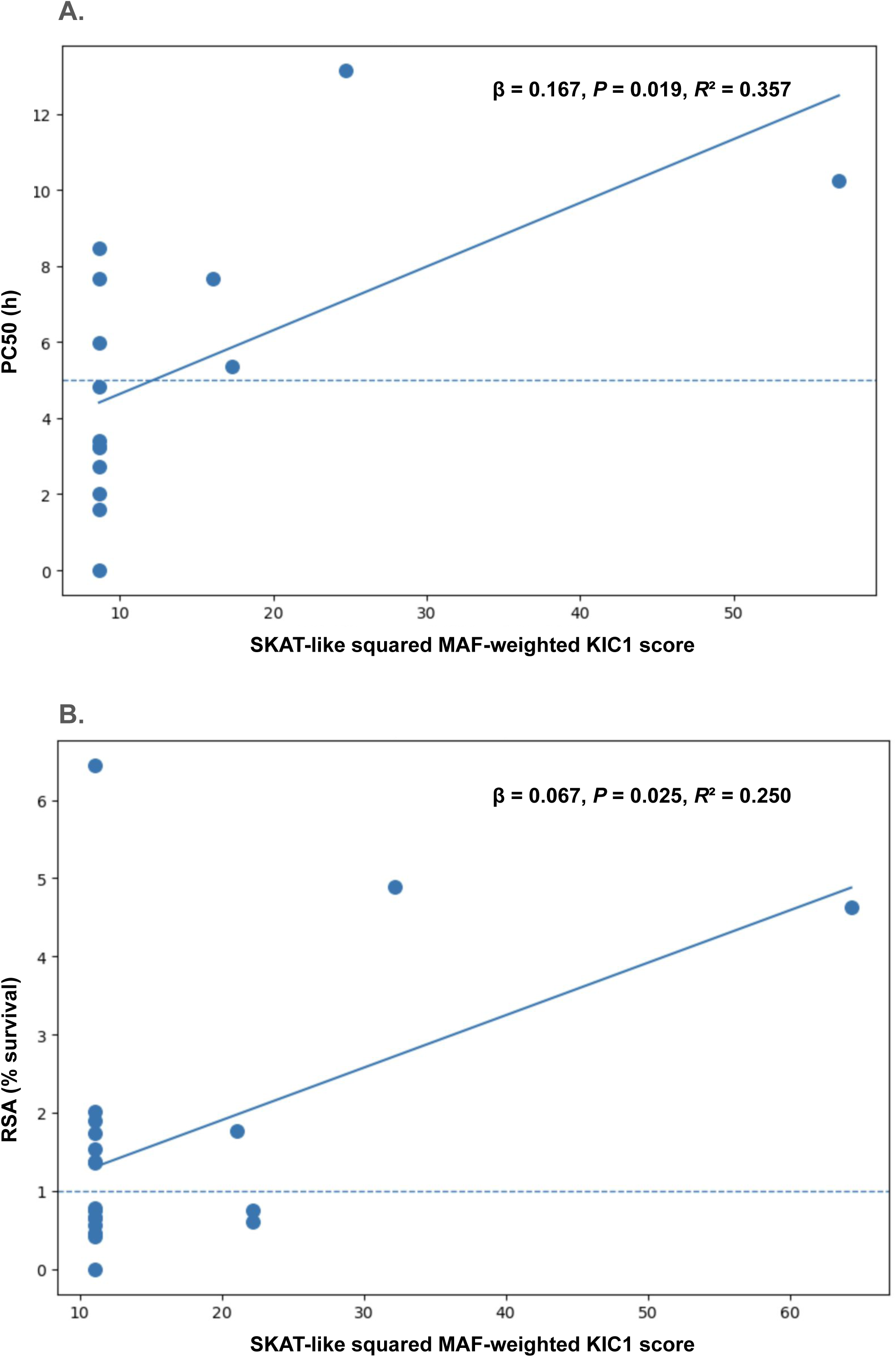
Rare-variant *pfkic1* gene score is associated with PC50 and RSA survival in CHT field isolates. **(A)** Relationship between the squared Madsen–Browning-weighted *pfkic1* variant score and PC_50_. Each point represents an individual parasite isolate. The solid line shows the fitted ordinary least-squares regression, which showed a positive association between the *pfkic1* score and PC_50_ (*P* = 1.86 × 10⁻², *R*² = 0.36). **(B)** Relationship between the squared Madsen–Browning-weighted *pfkic1* variant score and *in vitro* RSA survival. The solid line shows the fitted ordinary least-squares regression (*P* = 2.47 × 10⁻², *R*² = 0.25). The dashed horizontal line marks the 1% RSA survival threshold.

### Elastic net prioritizes partially overlapping *pfkic1* locus variants associated with PC_50_ and RSA survival

To identify individual *pfkic1* locus variants contributing to the gene-score associations, we fit Madsen–Browning-weighted elastic net regression models separately for PC_50_ and RSA survival. Leave-one-out cross-validation was used for hyperparameter selection, and bootstrap resampling was used to assess the stability with which each variant was retained. For PC_50_, the optimal model favored a sparse solution and retained four of seven *pfkic1* locus variants with non-zero coefficients. The intergenic variant Pf3D7_06_v3:249910: A>T had the largest model coefficient, while three missense variants at positions 251742, 251809, and 252876 (**Table S7**) were retained with smaller coefficients. For RSA survival, the optimal model retained four of seven variants, with a more distributed coefficient pattern. The missense variant Pf3D7_06_v3:253113: T>C showed the highest bootstrap selection stability for RSA and had a negative coefficient, while variants at 251742, 251809, and 249910 (**Table S7)** were also retained. Bootstrap stability selection over 1,000 resamples showed partial overlap between the PC_50_ and RSA models. The intergenic variant Pf3D7_06_v3:249910: A>T (**Table S7)** was retained in both models with moderate stability (π = 0.65 for PC_50_ and π = 0.65 for RSA). Missense variants at 251742 (N688Y) and 251809 (F665L) (**Table S7)** were also selected by both models with similar stability (π ≈ 0.66), suggesting a shared core signal at these positions. In contrast, the missense variant at 253113 (K231E) (**Table S7)** was most strongly prioritized in the RSA model (π = 0.77) but was not retained in the PC_50_ model, whereas the missense variant at 252876 (D310Y) (**Table S7)** was retained only in the PC_50_ model. These results suggest that the *pfkic1* locus association is not driven by a single variant but instead reflects a partially overlapping set of candidate variants associated with early clearance and *in vitro* RSA phenotypes.

### Exploratory genome-wide gene-score analysis nominates additional candidate loci associated with PC_50_ and RSA survival

Having tested the experimentally nominated *pfkic1* locus, we next applied the same Madsen–Browning-weighted gene-score framework genome-wide to ask whether additional loci showed nominal association with artemisinin-response variation in CHT field isolates. Gene-body variants and proximal intergenic variants within 50 kb were assigned to genes, and a weighted score was computed for each gene. Gene scores were tested against continuous PC_50_ and RSA survival, with group-based non-parametric comparisons used as supportive analyses. Benjamini–Hochberg FDR correction was applied to account for multiple testing, and QQ plots were used to assess the distribution and potential inflation of genome-wide gene-score associations (**Fig. S4A, B**). No genes reached genome-wide significance after FDR correction; therefore, these results were interpreted as exploratory nominal associations. Among the top-ranked nominal signals, GAMER (PF3D7_0805300) showed association with both PC_50_ and RSA survival and has previously been flagged in population-genetic analyses of ART-R [49]. Additional PC_50_-associated candidate loci included PfWD11 (PF3D7_1138800), a WD40-domain protein previously implicated in ART-R genetic-background analyses, and PF3D7_1462500, which was associated with both PC_50_ and RSA survival and lies near the chromosome 14 resistance-background region containing *pfarps10*. For RSA survival, top-ranked nominal loci included PF3D7_1434300, encoding the Hsp70/Hsp90 organizing protein homolog HOP, as well as VPS4, PTEX150, PK5, and an ApiAP2 transcription factor locus. Although these associations require validation in larger cohorts, they nominate candidate loci in pathways related to stress response, vesicle trafficking, exported-protein biology, transcriptional regulation, and cell-cycle control (**Fig. S5A, B**; **Tables S7 and S8)**.

## MATERIALS AND METHODS

### *Plasmodium falciparum in vitro* culture, maintenance and adaptation

Asexual-stage *Plasmodium falciparum* parasites were maintained in continuous culture at 4% hematocrit using O+ human erythrocytes (Biochemed Services, Winchester, VA; Interstate Blood Bank, Memphis, TN). Parasites were grown in complete medium consisting of RPMI 1640 supplemented with L-glutamine (Invitrogen), 50 mg/L hypoxanthine (Sigma-Aldrich), 25 mM HEPES (Sigma), 2 g/L D-glucose (Sigma), and 20 mg/L gentamicin (Gibco), along with 0.5% AlbuMAX II (Gibco) and 0.22% NaHCO₃ (Corning). Cultures were maintained under a controlled gas environment (5% CO₂, 5% O₂, 90% N₂) following standard conditions [50]. Culture adaptation of cryopreserved clinical isolates was performed as previously described [21], with minor modifications. Briefly, cryopreserved infected red blood cell samples were rapidly thawed at 37°C and recovered by stepwise washes using 12% NaCl, 1.6% NaCl, and 0.9% NaCl/0.2% dextrose. Recovered parasites were established in culture with fresh human O+ erythrocytes at 10% hematocrit in complete RPMI medium supplemented with Albumax, rather than pooled human AB+ serum. Cultures were maintained at 37°C in the malaria gas mix and monitored daily by Giemsa-stained thin blood smears, with daily medium changes until parasite growth was established and stocks were cryopreserved.

### Stepwise Selection of DHA-Resistant Lines

A recently adapted *P. falciparum* isolate from Bangladesh, CHT-S, carrying wild-type *pfkelch13* and exhibiting an RSA survival rate of <1%, was subjected to stepwise *in vitro* DHA selection. Two selection regimens were performed: 24-h DHA exposure in biological duplicate and 48-h DHA exposure in a single replicate. In parallel, the parental CHT-S line was maintained without DHA and designated CHT-S-sib. DHA concentrations were incrementally increased from 20 nM, 10× EC₅₀, to 1.6 µM, 800× EC₅₀, over 18 months. Following each DHA exposure, parasites were cultured drug-free until recovery to 1% parasitemia; the interval from the start of the DHA pulse to recovery defined one selection cycle. DHA concentrations were increased stepwise across successive selection cycles: 40, 60, 80, 100, 200, 400, 600, 800, 1200, and ultimately 1600 nM. During the early stages of selection, if parasites recovered within 12–14 days, the same DHA concentration was reapplied in the subsequent cycle; if recovery required more than 12 days, the next higher DHA concentration was applied. Three final resistant lines, CHT-R-24-1, CHT-R-24-2, and CHT-R-48, were derived, and RSA and post-DHA recovery assays were performed during selection to monitor the emergence of ART-R phenotypes.

### Plasmid construct for *pfkic1* -TGD

The TGD construct was created using pSLI-TGD plasmid (**Fig. S2A**) (Addgene: 1463000) following previously published protocols [51]. Briefly, primers were designed with NotI and MluI restriction sites: 5’-NotI-Stop-ATGless-KIC1-Fw (atagcggccgctaaACTAATGTTAATAATAATATGAACAATTCAGGC) and 3’-MluI-516-KIC1-Rev (ataacgcgtGGAATTATTAAAATTATCAGAGTTTCTTGTTTTTTC). The *pfkic1* fragment was amplified from CHT-S genomic DNA and cloned in the pSLI-TGD plasmid at the NotI/MluI sites.

### Parasite Transfection and Selections

Ring-stage parasites of CHT-S (5% parasitemia) were transfected with 50 µg of the final plasmid and selected with 5nM of WR99210 (Jacobus Pharmaceuticals). Once parasites were recovered, they were selected with neomycin (neo) (Geneticin, Fischer Scientific) at 400 µg/mL in three independent attempts. Neo selection was initiated with 1% parasitemia in 4% hematocrit for 10 days and waited till parasites were recovered, which were again treated with 5nM WR099210 for 4 cycles before preparing genomic DNA for integration check. Integration in the three selections were confirmed by PCR using 5’ integration primers (Int-up-Fw: gtgctgtgttttttaccttgtttttgtgtc and GFP-rev:GCATCACCTTCACCCTCTCCACTGACAG), 3’ integration primers (pARL-Fw:ggaattgtgagcggataacaatttcacacagg, and gene-int-Rev:GACAAATCCTTCATATTTGTATTATTATTTTCTGCG). The absence of the WT *pfkic1* locus was assessed with Int-up-Fw: gtgctgtgttttttaccttgtttttgtgtc and gene-int-Rev:GACAAATCCTTCATATTTGTATTATTATTTTCTGCG primers to ensure successful disruption.

### Ring Stage Survival Assay (RSA)

DHA selected and *in vitro* adapted parasites were phenotype for artemisinin susceptibility by ring-stage survival assay (RSA). RSAs were performed on 0-3h rings as previously described [21]. Briefly, parasites were tightly synchronized using two sorbitol treatments performed 40 h apart. Percoll-purified, late-stage segmented schizonts were then incubated with fresh RBCs for 3 h, followed by an additional sorbitol treatment. Cultures containing 0- to 3-h rings were adjusted to 2% hematocrit and 1% parasitemia and seeded in 24-well plates at 1 mL complete medium per well containing either 700 nM dihydroartemisinin (DHA) or 0.1% dimethyl sulfoxide (DMSO) as the vehicle control. After 6 h of incubation at 37°C, cultures were washed and returned to drug-free medium. Thin blood smears were prepared 66 h later, corresponding to 72 h after seeding, and parasite survival was assessed microscopically by quantifying next-generation viable rings with normal morphology. Parasitemia was estimated by counting 20,000 RBCs in treated and 10,000 in DMSO controls. Survival rates, expressed as percent parasites, were calculated by dividing the viable parasitemia in DHA-treated cultures by that in DMSO-treated controls and multiplying by 100 [16].

### Post DHA treated Recovery Assay

Cultures containing 0–3 h ring-stage parasites were diluted to a starting parasitemia of 1% and treated with 700 nM DHA for 6 h. After 6 h, DHA was washed off, and parasites were returned to drug-free media. Parasitemia was measured by microscopy beginning at 72 h post-treatment and continued at designated time points up to 192 h (8 days). Media changes were performed at 72, 120, and 168 h following DHA treatment. Parasitemia was estimated by microscopy by counting 10,000 RBCs.

### DNA extraction and Whole genome sequencing

*P. falciparum* cultures were expanded to ∼5% parasitemia, enriching for mature stages. Cultures were lysed using 0.05% (w/v) saponin, and parasite pellets were washed with 1× PBS. Following RNase treatment, genomic DNA was extracted using the QIAamp DNA Blood Mini Kit (Qiagen). DNA quantity and integrity were assessed using a NanoDrop spectrophotometer and Agilent TapeStation (gDNA assay). Between 1–5 µg of high-quality genomic DNA was used for downstream library preparation. Whole genome sequencing was performed similarly to our previously published study [43]. Briefly, genomic DNA were processed using the NEBNext Ultra II DNA Library Prep kit, with fragmentation performed by Covaris sonication to yield an average insert size of approximately 350 bp. Library quality and quantity were evaluated using KAPA HiFi qPCR, TapeStation DNA High Sensitivity assays, and Qubit High Sensitivity DNA measurements. Following equimolar pooling, the libraries were sequenced with 150 bp paired-end reads on an Illumina NextSeq 2000 P2 (300-cycle) flow cell at the Genomics and Bioinformatics Core Facility, University of Notre Dame. Base call files were converted to FASTQ format using Illumina’s onboard DRAGEN pipeline for downstream analysis.

### Read alignment and variant calling

The downstream analysis framework was adapted from MalariaGEN Pf7 [52]. Raw FASTQ files were adapter-trimmed using trimmomatic. The resulting high-quality reads were aligned to the *Plasmodium falciparum* 3D7 reference genome using BWA-MEM with default settings. SAM files were then converted to indexed BAM files using SAMtools. PCR duplicates were removed and base quality score recalibration was performed using the GATK pipeline. Variant calling was performed using GATK -HaplotypeCaller with sample_ploidy 1 and other default settings. The “*CombineGVCFs*” and “*GenotypeVCF*” commands were used to combine the individual vcf files and extract the genotypes. GATK “--selectVariant” command was used to extract the SNPs and indels and hard filtering was applied to remove low-quality SNPs (“QD < 2.0”, “FS > 60.0”, “MQ < 40.0”, “SOR > 4.0”, “MQRankSum < -12.5”, “ReadPosRankSum < -8.0”, “DP < 10”, “GQ < 10”) and biallelic SNPs were chosen to identify selection signatures from *in vitro* drug pressuring. Only the 21MB core genome region was used for downstream analysis after removing all hypervariable, centromeric, and subtelomeric regions.

### Variant Filtering and Selection Analysis

Genomic variants were identified from genotype VCF files generated for the three DHA-selected parasite lines along with corresponding sibling and parental controls. Variant processing was performed using a sequential filtering strategy. Initially, all variants (SNPs and indels) were retained, after which only biallelic SNPs were selected. Variants were then filtered based on read depth (DP) to remove low-confidence calls. Site-level quality control was applied by retaining only variants annotated with the “PASS” flag. Additional filtering was performed using genotype-level filter tags to exclude low-quality genotype calls. Following these steps, variants were restricted to the core genome to eliminate highly variable or repetitive regions. From the resulting high-confidence SNP set, unique variants in each 1600 nM DHA-selected line were identified by excluding variants present in both sibling and bulk controls. Functional annotation was used to classify variants, and non-synonymous SNPs (missense and stop-gain) were prioritized for downstream analysis. Genes harboring these variants were identified, and intergenic variants were analyzed separately. To identify convergent selection signals, genes containing non-synonymous SNPs in three independent 1600 lines were compared. Genes were retained if they contained distinct SNPs across lines and if these variants were absent in both sibling and parent controls. This approach enabled the identification of candidate genes under selection at the gene level rather than at specific nucleotide positions.

### Rare-variant gene-score analysis of the *pfkic1* locus and genome-wide variant burden

We quantified the contribution of *pfkic1* locus variation to RSA and PC_50_ by constructing a gene scoring test-based study [48]. Genotypes were called against the 3D7 reference genome; and for each SNP *i* and sample *j*, genotypes were converted into a binary indicator of alternate allele presence, where *A*ⱼᵢ = 1 if at least one alternate allele was observed and *A*ⱼᵢ = 0 otherwise; missing calls were encoded as 0. This yielded a binary genotype matrix A ∈ ℝⁿˣᵐ, where *n* denotes the number of samples and *m* the number of variants. For each retained non-synonymous (missense, stop-gained) or intergenic variant, *i*, the minor allele frequency (MAF) was computed as the mean of the corresponding column of A across all *n* samples. SNPs were assigned a Madsen-Browning weight *wᵢ* = 1 / √(MAFᵢ(1 − MAFᵢ)), with weights set to zero when MAFᵢ approached 0 or 1 [48]. To confer robustness to mixed allelic effect directions, we adopted a SKAT-like formulation [53] in which the genetic score was defined as *Sj* = Σ_i_ (*w*_i_ · *A*_ji_)², where the squared weighting allows contributions from variants acting in opposing directions to accumulate additively. Given the binary encoding of genotypes, the expression simplifies to *S*_j_ = Σ_i_ *w*^2^_i_ · *A*_ji_; PC_50_ and *in vitro* RSA survival values were used as primary phenotypes. Association between the gene score and each phenotype was tested using ordinary least squares (OLS) regression: Phenotype_j_ = β₀ + β₁ · *S*_j_ + ε_j_. Pearson and Spearman correlations were computed to characterize linear and monotonic associations, respectively. Group-wise comparisons in the context of KIC1 between PC_50_<5h and PC_50_>5h isolates and sensitive vs. resistant isolates were performed using the Mann–Whitney *U* test and Wilcoxon rank-sum test. To assess the stability of observed group differences in the KIC1 context, a permutation test was conducted by randomly shuffling gene scores across samples and recomputing the difference in group means over 10,000 iterations, from which an empirical *p*-value was obtained. For genomewide calculation using PC_50_ and RSA values as phenotypes, intergenic variants were assigned to the closest flanking gene for which an observed genomic interval could be estimated, provided that the distance from the SNP position to the nearest boundary of the gene’s observed interval did not exceed 50,000 bp [54,55]. Group-based analysis was performed using Welch’s two-sample t-test, Mann–Whitney U test, Wilcoxon rank-sum test, and Wald test p-values. Analyses were restricted to genes with at least two samples in each phenotype group. Multiple testing correction was applied to each set of p-values independently using the Benjamini–Hochberg false discovery rate (FDR) procedure. Genome-wide results were visualized using quantile–quantile (QQ) plots of observed versus expected −log₁₀(p) under the null, and Manhattan plots displaying −log₁₀(OLS p-values) across chromosomes ordered by genomic position. Significance thresholds corresponding to a Bonferroni-corrected α = 0.05 and a nominal α = 0.05 are indicated on Manhattan plots.

### Elastic net based penalized Regression with Stability Selection for Variant Prioritization

To localize the KIC1 signal to specific candidate variants, we fit Madsen-Browning-weighted elastic net regression models [56] with the matched phenotypes (PC_50_ and RSA, analyzed separately) as continuous outcomes and the per-SNP weighted genotype matrix as predictors. The elastic net estimator minimizes the penalized residual sum of squares:

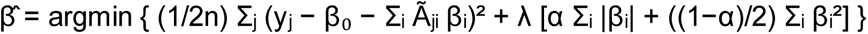

where λ controls the overall penalty strength and α ∈ [0, 1] interpolates between ridge (α = 0) and lasso (α = 1) regularization. The two hyperparameters were jointly selected by leave-one-out cross-validation over a grid of L1/L2 mixing ratios (α ∈ {0.1, 0.3, 0.5, 0.7, 0.9, 1.0}) and 100 automatically selected λ values per ratio, choosing the combination that minimized mean held-out prediction error. For each phenotype, *B* = 1000 bootstrap datasets were generated by sampling *n* isolates with replacement, 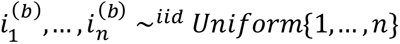, and fitting

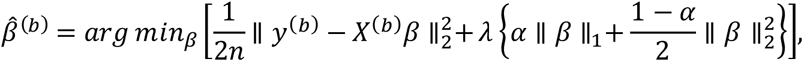

using the LOOCV-selected *λ* and *α*. Variant stability was defined 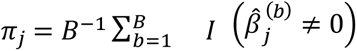, representing the proportion of bootstrap models selecting SNP *j*; variants with π_*j*_ ≥ 0.80 were considered robustly selected. Leave-one-out cross-validation was used in place of k-fold cross-validation given the limited sample size (n = 15 for PC_50_, n = 20 for RSA), where k-fold partitioning would yield highly variable fold-level error estimates. Models were fit using the ElasticNetCV and ElasticNet implementations in scikit-learn [57].

## DISCUSSION

While mutations in *kelch13* remain the dominant molecular markers of ART-R, laboratory evolution experiments and studies of natural parasite populations have demonstrated that resistance can emerge through multiple genetic routes [20,24]. Despite their genetic diversity, these determinants largely converge on two common functional strategies: reducing artemisinin activation through altered hemoglobin uptake and endocytosis [31,32], or enhancing parasite tolerance to artemisinin-induced cellular damage through mechanisms involving proteostasis, redox homeostasis, and stress-response pathways [20,24].

*In vitro* evolution has been a landmark approach for identifying ART-R determinants. Prolonged artemisinin selection of the F32-Tanzania line led to the discovery of *pfkelch13* as the major molecular marker of ART-R [17], and subsequent selection of a Malian parasite line identified an additional *pfkelch13* mutation sufficient to elevate RSA survival [58]. In parallel, selection experiments have revealed Kelch13-independent routes to reduced artemisinin susceptibility, most notably *pfcoronin* mutations in Senegalese parasites [59], stress-response and protein-damage pathways in 3D7 [60], and highly resistant field-isolate lineages with unconventional ring-stage phenotypes and no validated *pfkelch13* mutation [61].

Extending this approach to South Asian parasite backgrounds is important because ART-R in Bangladesh remains incompletely characterized and has not been linked to validated *pfkelch13* mutations [42,44,45]. In the CHTs, where most malaria transmission in Bangladesh is concentrated, our previous clinical and *in vitro* studies identified parasites with reduced artemisinin susceptibility despite wild-type *pfkelch13* [42,43]. We therefore applied long-term, stepwise DHA selection to the Bangladeshi CHT-S isolate to identify candidate determinants of ART-R in this regional genetic background (**Fig. 1**).

In our study, stepwise DHA pressure generated CHT-R lines with elevated survival of 0–3 h rings and progressively faster recovery after DHA exposure (**Fig. 1**). Notably, this phenotype emerged without acquisition of *pfkelch13* variants, distinguishing the CHT-R lines from Kelch13-driven laboratory-evolved ART-R system such as F32-Tanzania and the Malian selection line [17,58]. Thus, the CHT-S background provides a laboratory-evolved example of Kelch13-independent ART-R in which increased ring-stage survival is accompanied by improved post-drug recovery.

Whole-genome sequencing identified several candidate mutations across common genes in the three DHA-selected lines (**Table 1**). We prioritized *pfkic1* for functional validation because independent selections repeatedly acquired distinct predicted loss-of-function mutations in this gene, including two premature stop codons and a frameshift mutation (**Table 1**). This prioritization was further supported by the prior identification of KIC1 as the top-ranked Kelch13-interaction candidate in the original Kelch13 BioID study, placing it within the Kelch13-associated endocytic compartment [32]. The recurrence of *pfkic1* mutations in our selections resembles the DHA-selected Senegalese parasite lines in which independent selections converged on mutations in *pfcoronin*, a gene later shown to drive elevated RSA survival [59,62].

KIC1 has been identified as a component of the Kelch13-associated endocytic compartment, which includes AP-2μ, UBP1, Eps15, and other regulators of hemoglobin uptake and endocytosis [32,33]. Several components of this pathway, including AP-2μ, UBP1, and KIC7, have been implicated in reduced artemisinin susceptibility, supporting impaired endocytosis and reduced drug activation as a route to ART-R [35–37]. In our study, in the CHT-S background, independent disruption of *pfkic1* was sufficient to increase RSA survival and improve post-DHA recovery (**Fig. 2**), establishing KIC1 as a functional contributor to reduced artemisinin susceptibility in this Bangladeshi parasite background. The *kic1*-TGD lines showed higher RSA survival than the DHA-selected CHT-R lines (**Fig. 1C and Fig. 2A**), suggesting that complete genetic disruption of KIC1 may have a larger effect than the selected truncating mutations, or that additional compensatory adaptations evolved during long-term DHA pressure. In contrast, previous disruption of *pfkic1* or editing of African *pfkic1* natural variants in laboratory-adapted 3D7 did not produce an ART-R phenotype [35,63], highlighting the importance of parasite genetic background in shaping the consequences of perturbing the Kelch13 endocytic network. Although KIC1 has not previously been linked to ART-R, a piggyBac forward-genetic screen identified KIC1 as a gametocyte-development hit [64].

The relationship between KIC1 and Kelch13 in the CHT-S background may be more complex than a simple linear pathway. Although *pfkic1* disruption was readily obtained in the wild-type CHT-S parent, repeated attempts to disrupt *pfkic1* in *pfkelch13*-edited CHT-S lines were unsuccessful. This does not establish genetic incompatibility, but it suggests that combined perturbation of Kelch13 and KIC1 may impose a fitness cost in this background, consistent with their association with overlapping endocytic pathways.

The CHT clinical isolates provided a complementary view of artemisinin-response variation in natural Bangladeshi parasites. But whether the threshold determined by WHO [65] for GMS parasites appropriately capture resistance phenotypes in Bangladesh remains uncertain. Although these infections did not meet conventional delayed-clearance criteria [15,65] and lacked validated *pfkelch13* mutations, the lab adapted isolates showed heterogeneous RSA survival (**Fig. 3A**) and variable PC_50_ values (**Table 2 and** [42]). Notably, RSA survival was associated with PC_50_ but not with parasite clearance half-life (**Fig. 3B and 3C**), suggesting that early parasite reduction metrics may capture aspects of artemisinin response that are not reflected by the conventional half-life threshold. This distinction is relevant in light of recent analyses showing that RSA survival and parasite clearance half-life can be partially decoupled outside the classic Southeast Asian ART-R framework [66]. Together, these data suggest that CHT parasites harbor measurable variation in artemisinin response, even though they do not yet fit the canonical pattern of clinically delayed ART-R.

Beyond parasite clearance metrics, growing evidence indicates that artemisinin-response variation in natural populations can also arise through mechanisms not captured by validated *pfkelch13* markers alone. *pfkelch13* -wild-type have been reported in Asia and Africa, including Cambodian isolates with elevated RSA survival despite lacking *pfkelch13* mutations and Ugandan isolates with increased RSA survival linked to *falcipain-2a* rather than validated *pfkelch13* mutations [3,5,21,22,67]. Additional non-*pfkelch13* determinants, including *pfap2μ*, *pfubp1*, and *px1/PIN*, further implicate cytostomal/endocytic trafficking, hemoglobin uptake, and hemoglobin digestion as recurrent routes to altered artemisinin response [14,68,69].

After functionally validating KIC1 in the CHT-S background, we used the field-isolate dataset (**Table 2 and Fig. 3**) to ask whether this experimentally validated locus also showed evidence of association in naturally circulating parasites. Because individual *pfkic1* variants were rare and the isolate set was small, we interpreted the gene-score analysis as an exploratory locus-level test rather than a conventional variant-level association study. In this context, the association between the *pfkic1* score and artemisinin-response phenotypes provides supportive field-isolate evidence consistent with the experimental finding that perturbation of KIC1 can alter parasite response to DHA. The elastic-net analysis further suggested that the natural-isolate signal is unlikely to reflect a single causal SNP but may instead reflect a local haplotype or a combination of regulatory and coding variation. The upstream intergenic variant prioritized in both phenotype models raises the possibility that *pfkic1* expression, in addition to protein disruption, could modulate artemisinin response in field parasites. Notably, two coding variants prioritized in our models, KIC1 N688Y and KIC1 K231E (**Table S6)**, have also been reported among naturally occurring *P. falciparum* KIC1 variants in a recent study [63], suggesting presence of these overlapping variants in Bangladesh too. But the functional consequences of these missense variants in the CHT backgrounds remain unknown.

Extending the same gene-score framework genome-wide identified additional top-ranked nominal candidate loci beyond *pfkic1*. These signals remain exploratory given the limited sample size and lack of functional validation, but several mapped to pathways plausibly linked to artemisinin response. GAMER (PF3D7_0805300) showed the strongest cross-phenotype signal and has previously been flagged in population-genetic analyses of ART-R [49], while PfWD11 (PF3D7_1138800), a WD40-domain protein recently prioritized in population-genomic and transcriptomic analyses of the artemisinin-resistance genetic background, was associated with PC50 [70]. PF3D7_1462500 was associated with both PC50 and RSA survival and lies near the chromosome 14 resistance-background region containing *pfarps10*, raising the possibility that this signal reflects local haplotypic variation rather than a single causal gene [71]. Additional RSA-associated loci included HOP, VPS4, PTEX150, PK5, and an ApiAP2 transcription factor locus, collectively pointing to stress response, vesicle trafficking, exported-protein biology, transcriptional regulation, and cell-cycle control as candidate pathways that may contribute to artemisinin-response variation in CHT parasites.

Together, these findings support a model in which artemisinin-response variation in CHT parasites can arise through genetic routes beyond canonical *pfkelch13* mutations. The functional validation of KIC1 establishes one experimentally defined route to reduced artemisinin susceptibility in a Bangladeshi parasite background, while the field-isolate analyses suggest that natural variation at *pfkic1* and other candidate loci may also shape local differences in parasite response. Future work testing whether natural *pfkic1* variants alter KIC1 expression or function, and whether the specific genome-wide candidate loci identified here modify artemisinin response in CHT parasites, will help determine how these known resistance-associated pathways operate in this regional parasite population. Overall, this study shows that experimental evolution in a locally relevant parasite background can uncover region-specific determinants of reduced artemisinin susceptibility that may complement *pfkelch13*-focused molecular surveillance.

## FUNDING INFORMATION AND ACKNOWLEDGEMENTS

This work was supported by the National Institutes of Health, NIAID (#1R21AI180663-01A1 to A.M.; 2024–2026), Indiana Clinical and Translational Sciences Institute (ICTSI) to A.M. (2023–2025), and the University of Notre Dame, College of Science, Center for Rare and Neglected Diseases. M.K.N. was supported by a Ph.D. fellowship from the Eck Institute of Global Health and College of Science (University of Notre Dame). We thank Dr. Kasturi Haldar for establishing the clinical capacity in the CHTs. We thank the Genomics and Bioinformatics Core Facility of Notre Dame for the whole-genome sequencing.

## DATA AND CODE AVAILABILITY

The codes are available in this github link https://github.com/NirjharBhattacharyya/Bangladesh_KIC1/tree/main.

