## Supplementary material for "Experimental evolution and functional genetics identify KIC1 as a determinant of reduced artemisinin susceptibility in Bangladeshi *Plasmodium falciparum*": Fig S1-S5

**Supplementary Figs**

**Fig. S1**. SNP composition in the in vitro selected lines. A: SNP (left) and B: INDEL (right)composition in CHT-S, CHT-S-sib and in vitro DHA selected lines. C. SNP composition within genes and in intergenic regions.

**A.	B.**

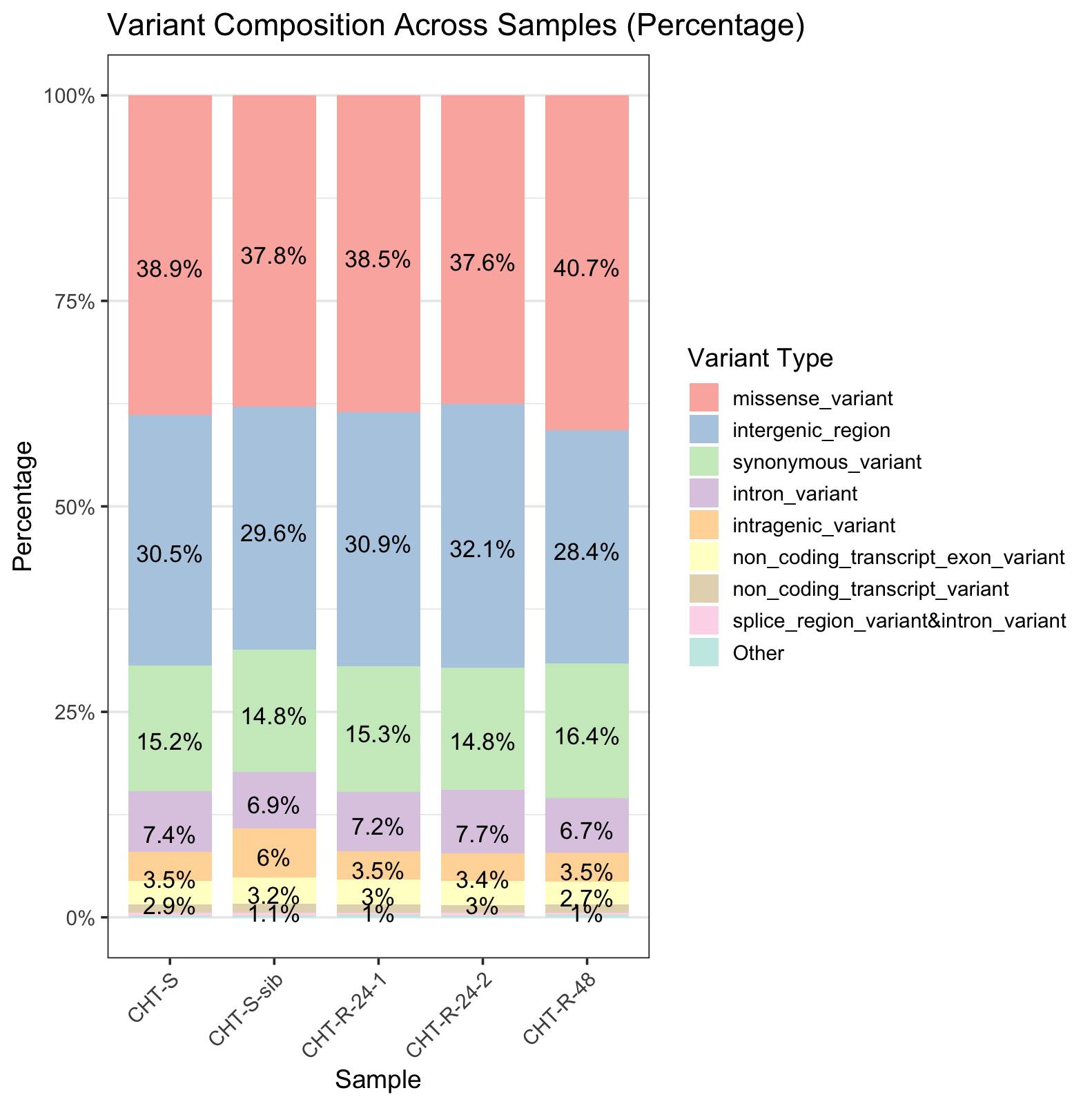

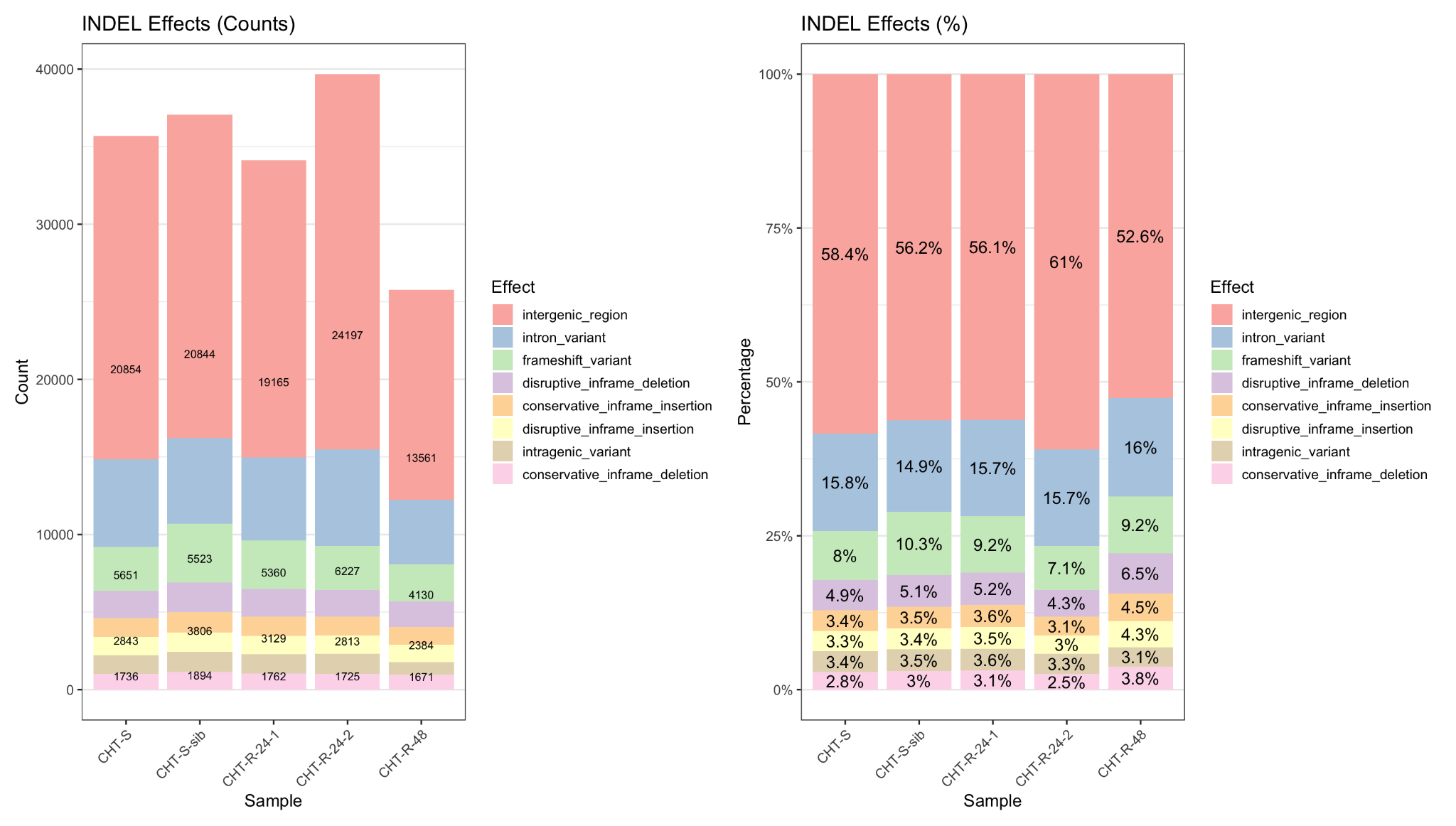

**C.**

**CHT-R-48	CHT-R-24-2	CHT-R-24-1**

**Fig. S2**. A. Schematic of the SLI-TGD strategy. B. Agarose gel pictures of the PCR integration tests for the successful kic1 disruption. The presence of amplicons in the 5' and 3' integration lanes and the absence of the WT allele confirm that the gene has been successfully disrupted in TGD-1, TGD-2 and TGD-3.

A

B

**Fig. S3**. Distribution of SKAT-like squared MAF-weighted KIC1 scores across artemisinin-response groups. Violin plots show score distributions, boxplots indicate the median and interquartile range, and points represent individual parasite strains. Strains were grouped by PC50 <5 h versus PC50 ≥5 h (A) and by RSA <1% sensitive versus RSA ≥1% resistant (B). Differences between RSA groups were evaluated using Welch’s *t*-test and the Mann–Whitney test.

**A.**

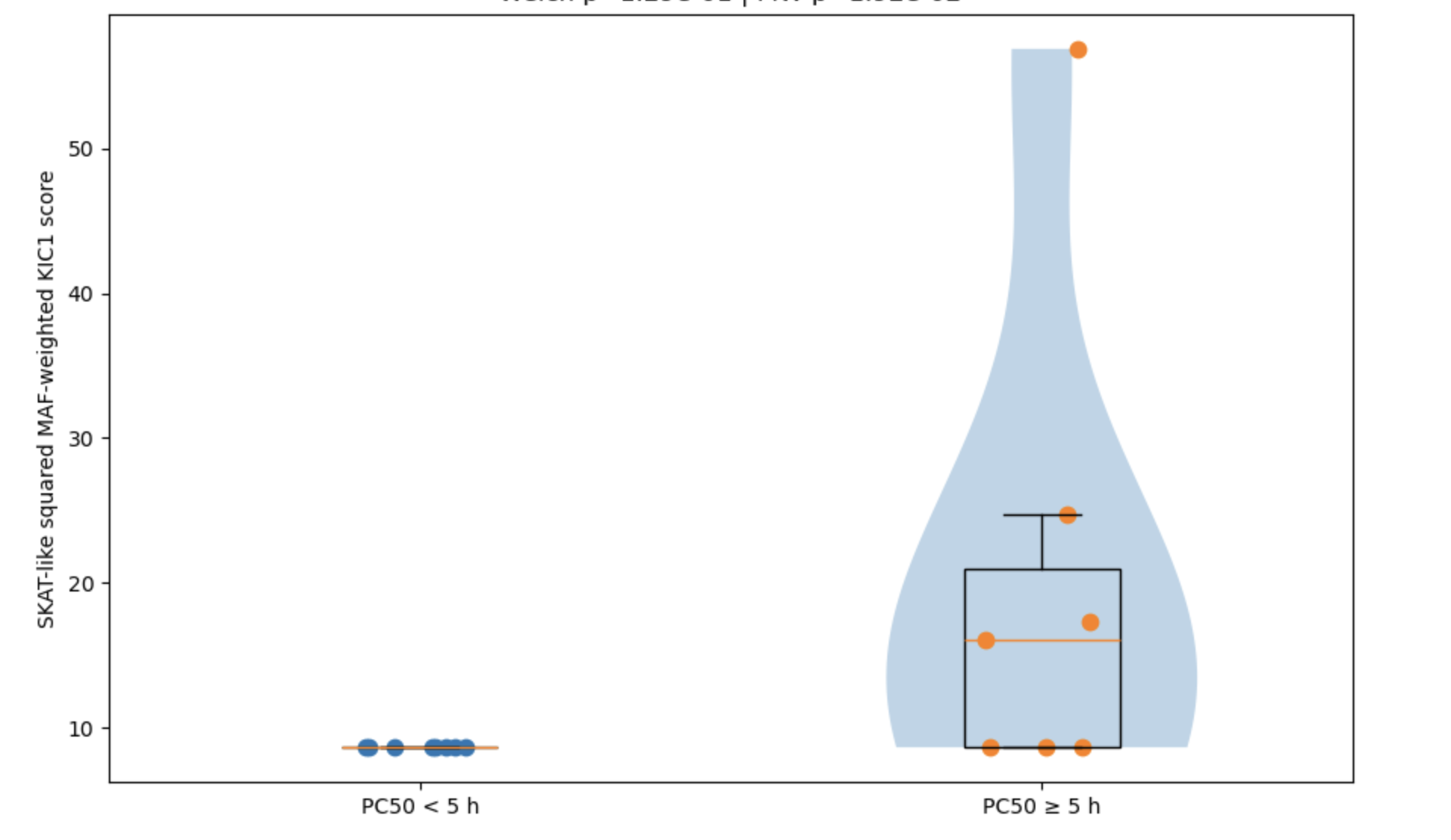

**B.**

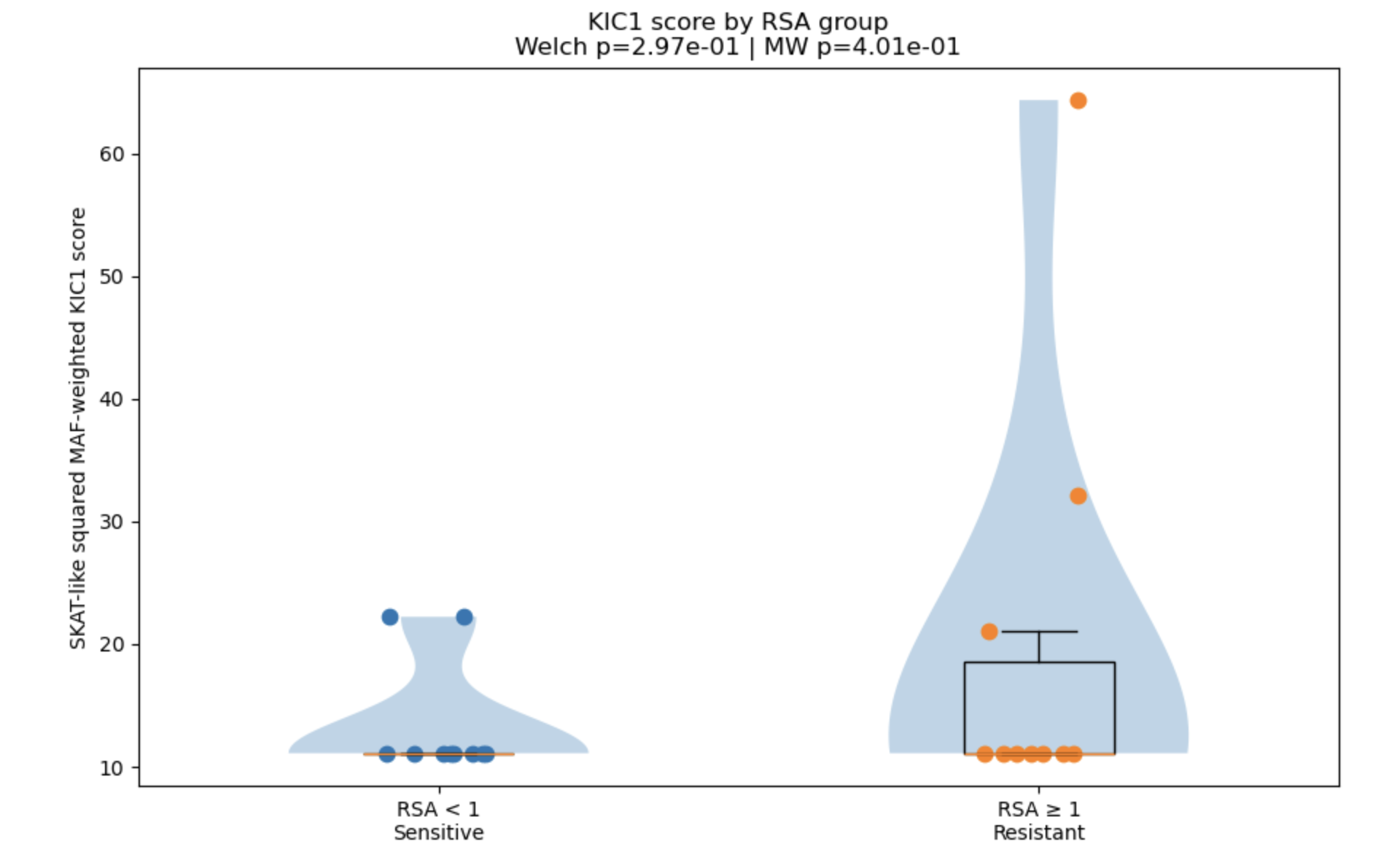

**Fig. S4. Gene-level association results for RSA and PC50 phenotypes.** Quantile–quantile plots compare observed and expected −log_10_(p) values from gene-score OLS analyses after excluding synonymous variants for RSA (A) and PC50 (B).

**A. B.**

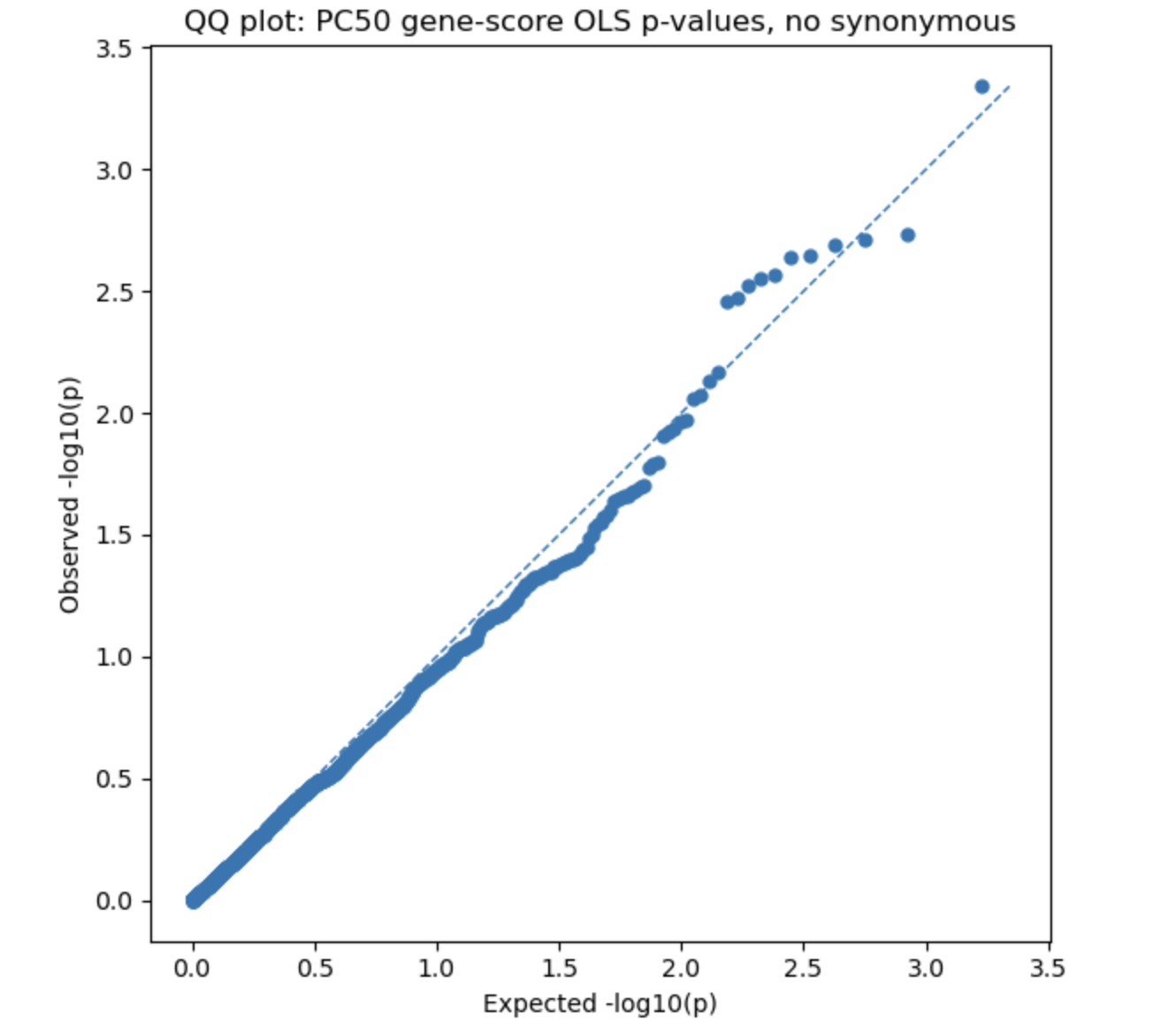

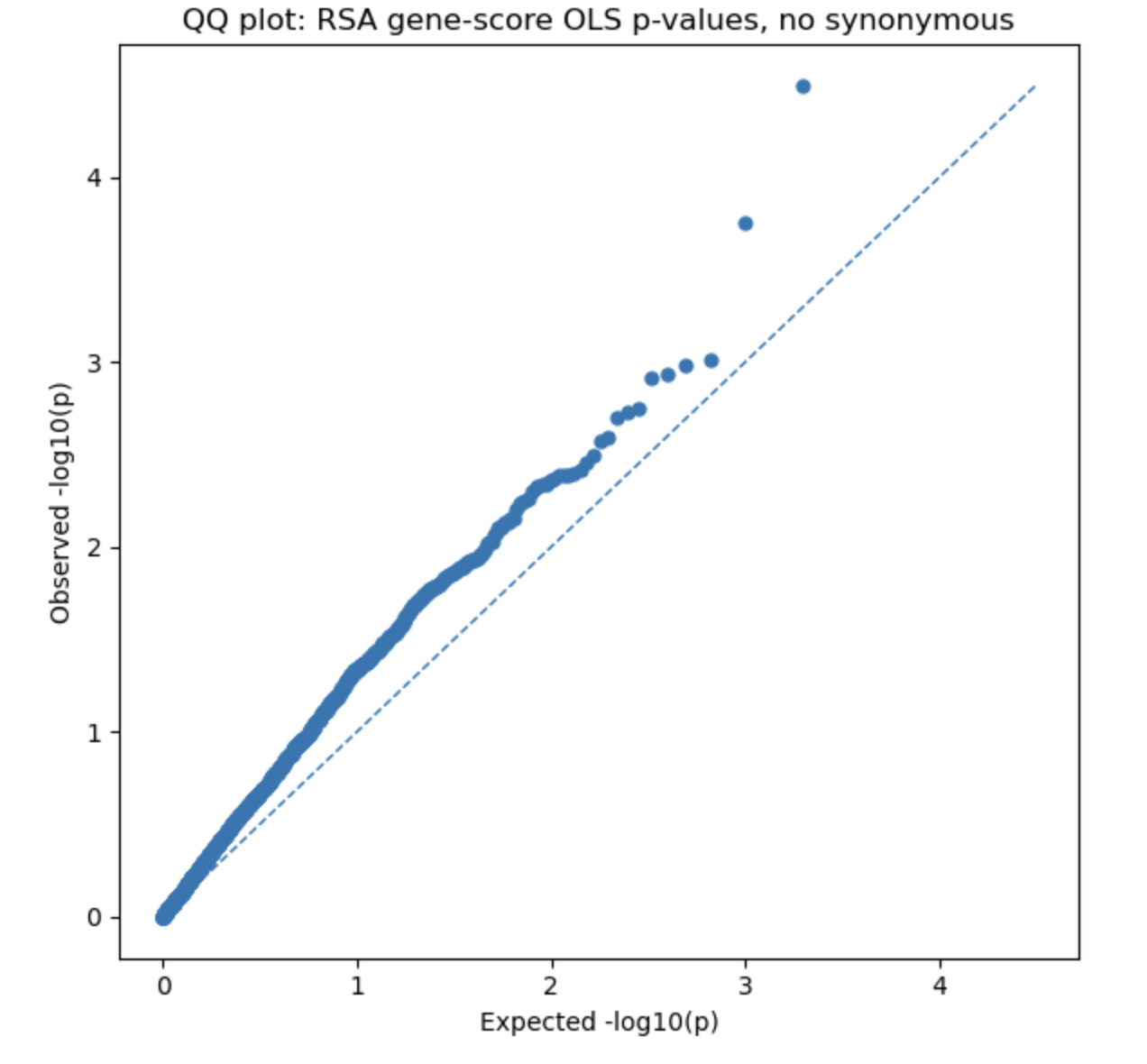

**Fig. S5. Genome-wide gene-score analysis identifies nominal candidate loci associated with PC50 and RSA survival in CHT field isolates.**

**(A)** Genome-wide Manhattan plot showing gene-level associations with PC_50_. **(B)** Genome-wide Manhattan plot showing gene-level associations with *in vitro* RSA survival. Each point represents a gene-level Madsen–Browning-weighted score positioned by chromosomal location. The y-axis shows the −log₁₀-transformed OLS association *P* value. The lower dashed line indicates the nominal significance threshold (*P* = 0.05), and the upper dashed line indicates the Bonferroni-corrected genome-wide significance threshold. Selected top-ranked nominal candidate genes are labeled.

**A**

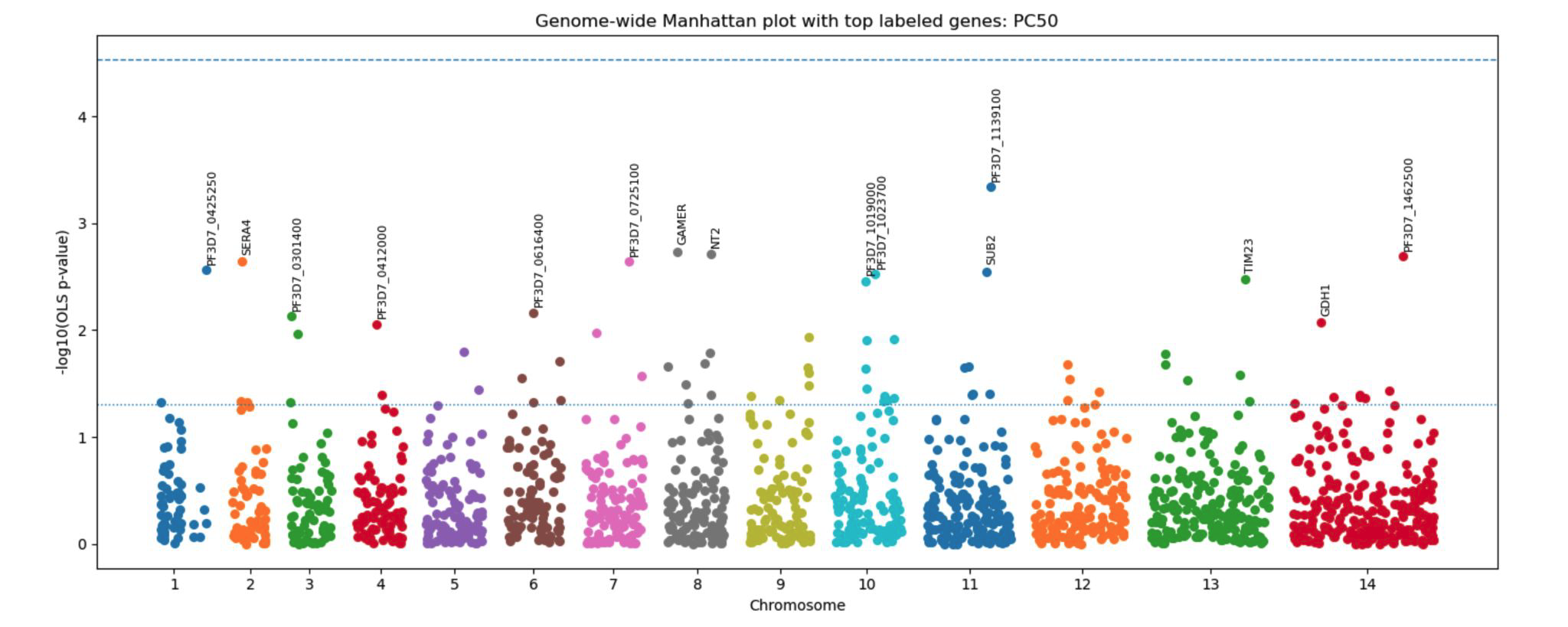

**B**
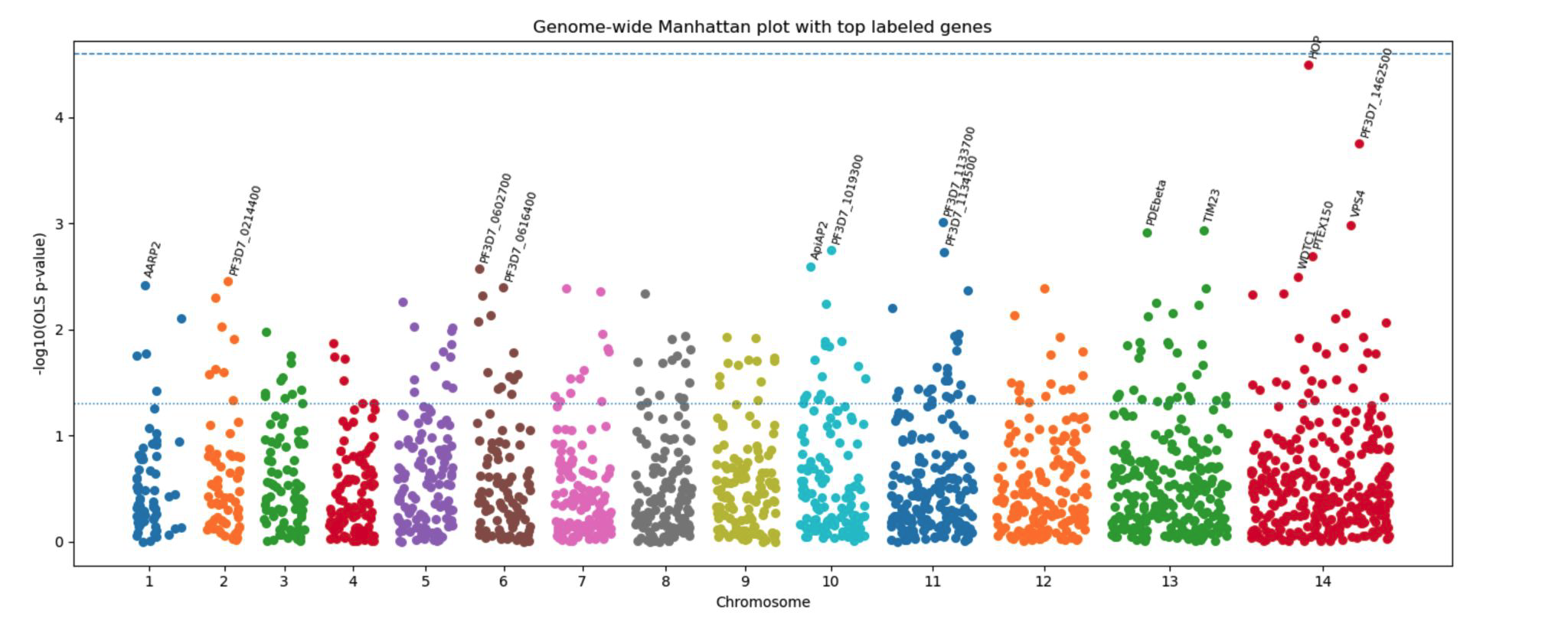
